# Comparative phylogeography reveals how a barrier filters and structures taxa in North American warm deserts

**DOI:** 10.1101/2020.06.17.157842

**Authors:** Kaiya L. Provost, Edward A. Myers, Brian Tilston Smith

**Affiliations:** Department of Ornithology, American Museum of Natural History, New York, New York; Richard Gilder Graduate School, American Museum of Natural History, New York, New York; Department of Vertebrate Zoology, National Museum of Natural History, Washington, District of Columbia

**Keywords:** biogeographic barrier, comparative phylogeography, functional traits, genetic diversity, isolation-by-distance, isolation-with-migration, neural net

## Abstract

The study of biogeographic barriers have been instrumental in understanding the evolution and distribution of taxa. Now with the increased availability of empirical datasets, it is possible to infer emergent patterns from communities by synthesizing how barriers filter and structure populations across species. We assemble phylogeographic data for a barrier and perform spatially-explicit simulations to quantify temporal and spatial patterns of divergence, the influence of species traits on these patterns, and understand the statistical power of differentiating alternative diversification modes. We incorporate published datasets to examine taxa around the Cochise Filter Barrier, separating the Sonoran and Chihuahuan deserts of North America, to synthesize phylogeographic structuring across the community with respect to organismal functional traits. We then use a simulation and machine learning pipeline to assess the power of phylogeographic model selection. Taxa distributed across the Cochise Filter Barrier show heterogeneous responses to the barrier in levels of gene flow, phylogeographic structure, divergence timing, barrier width, and divergence mechanism. These responses vary concordantly with locomotor and thermoregulatory traits. Many taxa show a Pleistocene population genetic break, often with introgression after divergence. Allopatric isolation and isolation-by-environment are the primary mechanisms purported to structure taxa. Simulations reveal that in spatially-explicit isolation-with-migration models across the barrier, age of divergence, presence of gene flow, and presence of isolation-by-distance can confound the interpretation of evolutionary history and model selection by producing easily-confusable results. By synthesizing phylogeographic data for the Cochise Filter Barrier we show a pattern where barriers interact with species traits to differentiate taxa in communities over millions of years. Identifying the modes of differentiation across the barriers for these taxa remains challenging because commonly invoked demographic models may not be identifiable across a range of likely parameter space.

## Introduction

Biogeographic barriers, which separate taxa and restrict gene flow via unsuitable habitat or impeded dispersal, have shaped the fields of evolutionary biology and ecology (e.g., Simpson, 1940; Lomolino, Sax, and Brown, 2004). Observing spatially disjunct taxa led to development of theory in species concepts (e.g., Mayr, 1942; de Queiroz, 2005), island biogeography (e.g., Janzen, 1967; MacArthur and Wilson, 1967), hybridization (e.g., Hoskin, Higgie, McDonald, and Moritz, 2005), and taxonomy (e.g., Helbig, Knox, Parkin, Sangster, and Collinson, 2002). The unifying factor across these disciplines is that the barrier itself is directly or indirectly linked to the process that generated the pattern of interest. Understanding the mechanisms that produce these patterns has been a major focus in historical biogeography, and efforts have expanded with the increasing number and complexity of phylogeographic studies (see Garrick et al., 2015).

Phylogeography has the dual aim of characterizing how diversity is distributed across the landscape and proposing the temporal framework for when the diversity formed (Avise et al., 1987). In addition, phylogeographic approaches aim to infer the processes that have promoted genetic diversity across the landscape (Avise, Bowen, and Ayala, 2016). Early studies provided insight into the distribution of genetic variation within species and species complexes that often coincided with geographic features or environmental gradients (e.g., Taberlet and Bouvet, 1994; Kozak, Blaine, and Larson, 2005). The extension of phylogeographic approaches with advances of molecular dating yielded insight into the timing of divergence between populations (e.g., Fouquet et al., 2010; Matos et al., 2016). Further advancements in DNA sequencing technology and statistical modeling allowed researchers better identify the patterns and processes associated with biogeographic barriers (Hickerson et al., 2010). However, the study of species diversification across barriers has been hampered by conceptual and logistical challenges. One such challenge is that although many taxa might show phylogeographic structure that is concordant with a barrier, this does not indicate that the barrier itself caused the genetic differentiation. For example, processes such as speciation over environmental gradients (Nosil, 2008; but see Bierne, Gagnaire, and David, 2013), behavioral selection (Zhang et al., 2012), or abundance troughs (Barton and Hewitt, 1981; Barrowclough, Groth, Mertz, and Gutierrez, 2005) could all produce similar patterns.

A second challenge is that a barrier may cause differentiation in some taxa, but it may not affect the entire biota at once; this is known as a barrier being semi-permeable or having filters (Simpson, 1940). Filters are hypothesized to be mediated by organismal traits such as morphology (Dick, Hardy, Jones, and Petit, 2008; Hanson, Fuhrman, Horner-Devine, and Martiny, 2012), reproductive isolation (Christe et al., 2016; Provost, Mauck, and Smith, 2018), physiology and niche (Castoe, Spencer, and Parksinson, 2007; Edwards, Lloyd, and Armbruster 2018), or population demographics and drift (Carnicer, Brotons, Stefanescu, and Penuelas, 2012). The mechanisms potentially underlying filter barriers are numerous, possibly barrier- and taxon-specific, and likely not mutually exclusive. Identifying which traits are important for causing isolation requires knowing which taxa are and are not separated at a barrier, where the barrier is relative to phylogeographic breaks between populations, and what functional traits are correlated with differentiation across the filter. To illuminate the actual role of filter barriers in diversification, and the mechanisms causing it, an understanding of the processes of differentiation and variation in the context of communities is required.

Advances in demographic modeling has enabled phylogeographers to distinguish between alternative modes of population differentiation (e.g., Gutenkunst, Hernandez, Williamson, and Bustamante, 2009; Gravel, 2012). Many of these approaches must manage computation expense while maximizing the information used (e.g., full likelihood methods, Bayesian methods) or alternately running quickly using approximate methods but requiring large amounts of summary statistics that may not use all of the data collected (e.g., ABC; Hickerson, Stahl, and Lessios, 2006; Jackson, Morales, Carstens, and O’Meara, 2017). New approaches allow testing demographic models on multiple taxa simultaneously (Satler and Carstens, 2017; Xue and Hickerson, 2017) and incorporating species abundances (Overcast, Emerson, and Hickerson, 2019). A common outcome of demographic analyses is overwhelming support for isolation-with-migration models (Nosil, 2008; but see Cruickshank and Hahn, 2014), which has two important implications. First, support for isolation-with-migration over pure allopatry challenges the view that differentiation is a discrete event. Second, filtering dynamics of barriers are likely more complex than recognized if gene flow occurs across them. Gene flow between diverging populations erodes the signal of that divergence by homogenizing the genome (see Eckert and Carstens, 2008), so estimating whether it has occurred is essential in detecting if divergence originally happened. Though detecting the timing and magnitude of gene flow is critical, using demographic model selection to do this can be difficult as models can often be indistinguishable (Roux et al., 2016). Further, most models assume that populations are panmictic, despite the prevalence of isolation-by-distance (IBD), and violating the assumptions of the model may compromise inference (Battey, Ralph, and Kern, 2020). Accounting for IBD in models is now relatively straightforward using the suite of recently developed spatially-explicit methods (e.g., Bradburd, Coop, and Ralph, 2018; Currat, Arenas, Quilodràn, Excoffier, and Ray, 2019), especially when combined with sophisticated statistical analyses like machine learning. Machine learning approaches relieve some of the statistical and computational burdens of demographic model selection by processing large amounts of data unconstrained by assumptions like data normality (Ripley, 1996). One type of machine learning that is particularly powerful for but underused in evolutionary biology is supervised machine learning, because it can incorporate what is already known about the data when making inferences (Schrider and Kern, 2018). Combined with simulation software developed over the past decades (e.g., Hickerson, Stahl, and Takebayashi, 2007; Gutenkunst et al., 2009), the phylogeographer has a powerful toolkit for estimating the evolutionary histories of their study groups.

Here, we use an exemplar biogeographic system, the Cochise Filter Barrier (CFB; Fig. 1), to explore the community-wide impact of a barrier on genetic differentiation within species, which traits mediate phylogeographic structure, and what analytical tools have been used to estimate those impacts. The CFB is a well-studied environmental and physical barrier that corresponds with a known transition zone between two biotas (Remington, 1968; Swenson and Howard, 2005) that divides the Sonoran and Chihuahuan Deserts of North America. Dynamic geological and ecological factors created this barrier, though the timing of the formation of the deserts is disputed (Wilson and Pitts, 2010). The southern region of the CFB dates back to the uplift of the Sierra Madre Occidental during the Oligo-Miocene, whereas the northern formed during the Plio-Pleistocene glacial cycles (Van Devender, 1990; Holmgren, Norris, and Betancourt, 2007) and the uplift of the Colorado Plateau (Spencer, 1996). In the north, xeric habitat connecting the deserts was ephemeral during the Plio-Pleistocene, with glacial advances repeatedly replacing aridlands with forests (Thompson and Anderson, 2000; Hafner and Riddle, 2011). This likely created desert refugia east and west of the CFB. Environmental gradients also exist between the hotter, wetter Sonoran Desert and the cooler, dryer Chihuahuan Desert, including elevational and vegetational differences and a high altitude grassland plain between the two (Shreve, 1942; Reynolds, Kemp, Ogle, and Fernández, 2004; Figure 1). Interactions between the environment, landscape, and organisms gave rise to the region’s biodiversity; some taxa are continuously distributed with no evidence of differentiation, while others show strong phylogeographic structure across the CFB (e.g., Zink, Kessen, Line, and Blackwell-Rago, 2001; Riddle and Hafner, 2006). The barrier is not one sharp break; rather, the biota is thought to turn over approximately between 112–108°W longitude (Pyron and Burbrink, 2010; Hafner and Riddle, 2011). The barrier is also associated with the Western Continental Divide (Castoe et al., 2007). Despite this, there is no quantitative estimate of the position and width of the barrier across taxa. Without understanding this, it is difficult to infer whether taxa have all been impacted by the same filter and which organismal traits may have been selectively advantageous for crossing the barrier and maintaining gene flow. It is also unclear when divergence happened, with estimates of population divergence ranging from the Miocene through the Pleistocene (e.g., Myers, Hickerson, and Burbrink, 2017), or whether dispersal occurred after lineage formation. Knowing when and where taxa diversify can lead to inferences on the processes that cause the initial divergence between populations, as well as further insights on post-divergence gene flow and the impact of IBD within structured lineages.

**Figure 1:**
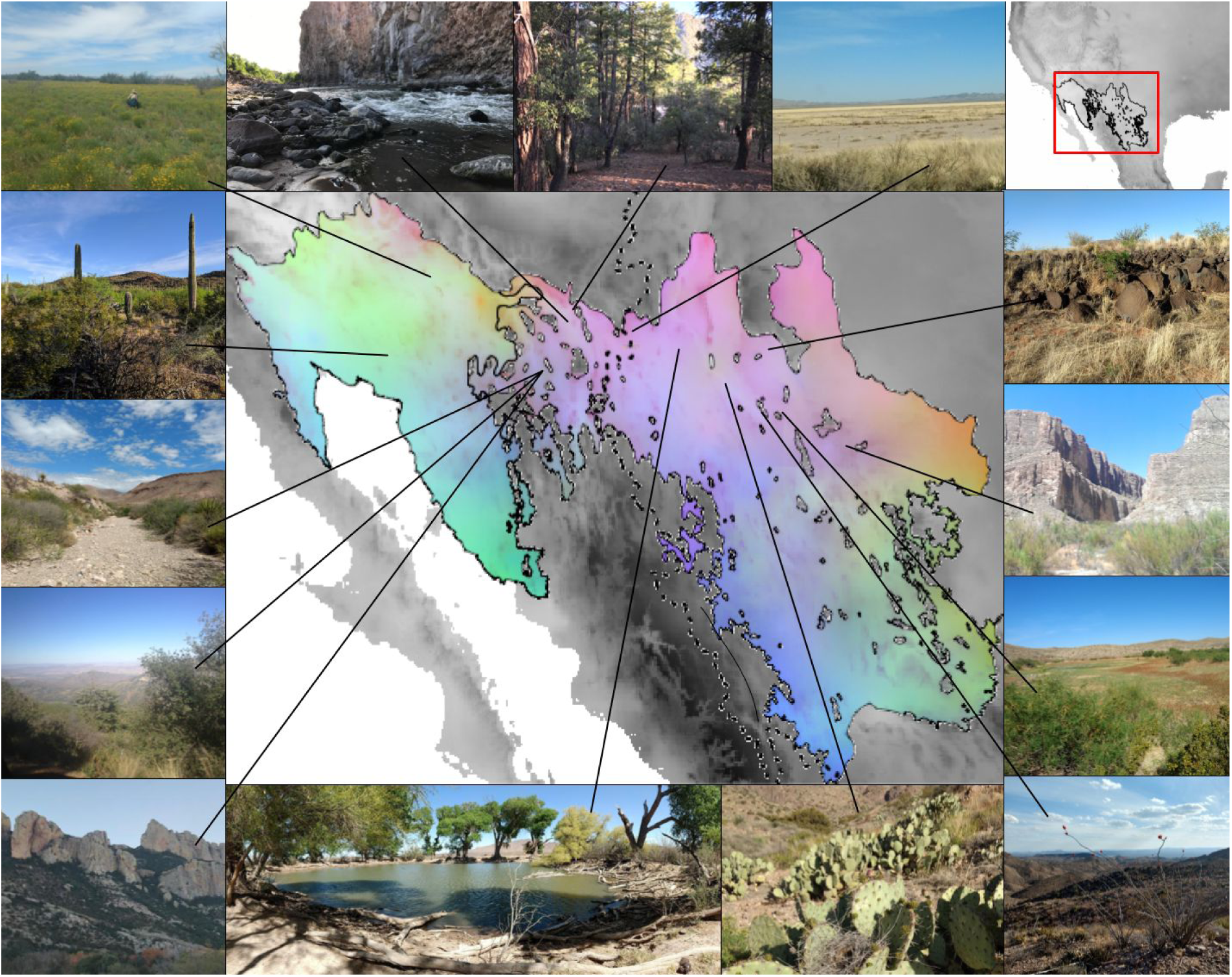
Examples of habitat and climatic variation in the Sonoran and Chihuahuan deserts. Center shows Sonoran and Chihuahuan desert outlines (Olson et al., 2001) and dotted line indicates the Western Continental Divide. Colors represent results of a PCA of climatic variables (see Appendix 1), with the first three principal components mapped to green, red, and blue channels of the image, respectively. More green indicates hotter summer temperatures. More red indicates more variable temperatures across the year. More blue indicates more summer precipitation. Areas outside of the deserts are the same principal components converted to grayscale. Top right shows the section of North America that the center region displays. Photographs around the map show representative habitats across the region, with a line leading to the point on the map where the photograph was taken.

We describe the diversity in genetic structure in taxa across the CFB. We synthesize what phylogeographic analyses have been used to understand the system (e.g., divergence dating, estimates of gene flow), which taxa are separated at the CFB, where the separation is, and when divergence occurred. We ask whether the location, age, and width of the barrier are consistent across taxa, because discrepancies can illuminate the filtering mechanisms that structure organisms. Further, we ask if variation in functional traits corresponds with the presence of phylogeographic structure. Under a dispersal limitation hypothesis, the vagility of organisms should be correlated with how much phylogeographic structure they display. A barrier should have a stronger impact on less vagile organisms. Under a thermoregulatory hypothesis, we expect that the ability of an organism to regulate their body temperature should be correlated with phylogeographic structure. Ectotherms may be more affected by environmental filters and more likely to experience reduced gene flow in the face of spatiotemporal climatic changes. We simulate genomic data under competing diversification scenarios across the CFB to determine whether genomic datasets and demographic models can further clarify how diversification proceeded. We aim to describe how synthesizing across empirical, theoretical, and simulated phylogeographic studies can lead to novel insights about the mechanisms of divergence across biogeographic barriers.

## Methods

### Characterizing phylogeographic breaks across the CFB

Understanding the filtering mechanisms operating on taxa distributed across barriers requires understanding where phylogeographic breaks are and which taxa are impacted. To quantify the patterns and frequency of modes of divergence across the CFB, we synthesize genetic patterns from published phylogeographic studies with inter- and intraspecific sampling across the CFB. We recorded the publication date of each study, data type (e.g., single-locus, SNP, etc.), and number of loci. Finally, we characterized each taxon based on their type of locomotion (following Burbrink et al., 2016) and type of thermoregulation (endo-vs ectothermic) to assess whether these traits interacted with the filter to impact the degree of population structuring. We also quantified elevational preference (lowland, montane, or both) to control for elevational differences on diversification. To assess the effects of traits on divergence times, we performed generalized linear mixed modeling (GLMMs) via the ‘lme4’ package version 1.1-21 (Bates, Mächler, Bolker, and Walker, 2015) in R version 3.6.1 (R Core Team, 2019) with and without accounting for taxonomy and elevation (see Appendix 1 for supplementary analyses).

For each taxon, we examined if a specific mode of diversification across the barrier was implicated by the authors of the respective studies (i.e., allopatry, ecological, hybridization, sexual selection, or polyploidy). Second, we assessed whether there was phylogeographic structure across the CFB, when structure arose, and if populations were reciprocally monophyletic with respect to individual loci. Because of the ambiguity in the placement of the CFB we chose to characterize whether there was a phylogeographic break between 118–99°W longitude, a much larger range than previously thought (Pyron and Burbrink, 2010; Hafner and Riddle, 2011; Myers et al., 2019). In addition, we assessed the width of the contact zone for each taxon by taking the localities of specimens used in each study, or if specimen information was not available, by assessing placement of specimens on figures. We ignored other biogeographic breaks known to occur within the species’ ranges. Missing data for variables not estimated for a given taxon were coded as ambiguous.

To determine when phylogeographic structure formed, we categorized the divergence time between the desert populations by epoch: Pleistocene (2.58 Ma to 11.7 Ka), Pliocene (5.33 Ma to 2.58 Ma), Miocene (23.0 Ma to to 5.33 Ma), or overlapping two epochs. We separated dates by epoch to allow broad comparisons in divergence across the community. Some studies suggested epochs of divergence between populations without explicit estimates of the divergence date, which we noted. When applicable, we included the full range of error around an estimate. In addition, to assess whether taxa were monophyletic with respect to the CFB, we visually assessed gene trees present in the study. Finally, we assessed whether gene flow across the barrier was explicitly estimated. If gene flow was not explicitly estimated, we considered individuals who show admixture in clustering analyses as a proxy for gene flow; we recognize that a high coefficient of ancestry could result from incomplete lineage sorting of ancestral polymorphisms. We then recorded whether gene flow was present, absent, or ambiguous.

### Simulation of discrete population structure and IBD

From our assessment of the literature across the CFB, two patterns were consistently found: discrete phylogeographic structure, or IBD without a discrete transition between populations. However, it is possible that both of these patterns are simultaneously observable in empirical population genetic data (Bradburd et al., 2018) and could be distinguished using a simulation-based approach. We used the program SLiM 3.1 (Haller and Messer, 2019) to build models that explored how the processes of allopatric isolation and speciation-with-gene-flow interact to generate divergence across a biogeographic barrier.

First, we simulated four demographic models, which varied by genetic structure and introgression: 1) a single population model, in which only one panmictic population exists across the entire region, 2) a pure isolation model, in which two populations exist but no gene flow is allowed between them, 3) an isolation-with-migration model, as in pure isolation but allowing gene flow continuously between the populations at a low rate, and 4) a secondary contact model, with gene flow allowed during the final 1,000 generations of the simulation (Figure 2). We chose 1,000 generations of the simulation to represent late-Holocene secondary contact. These four models represent different underlying processes of diversification that are commonly tested for in phylogeographic studies, and with them we evaluated how gene flow and divergence across the CFB varied.

**Figure 2:**
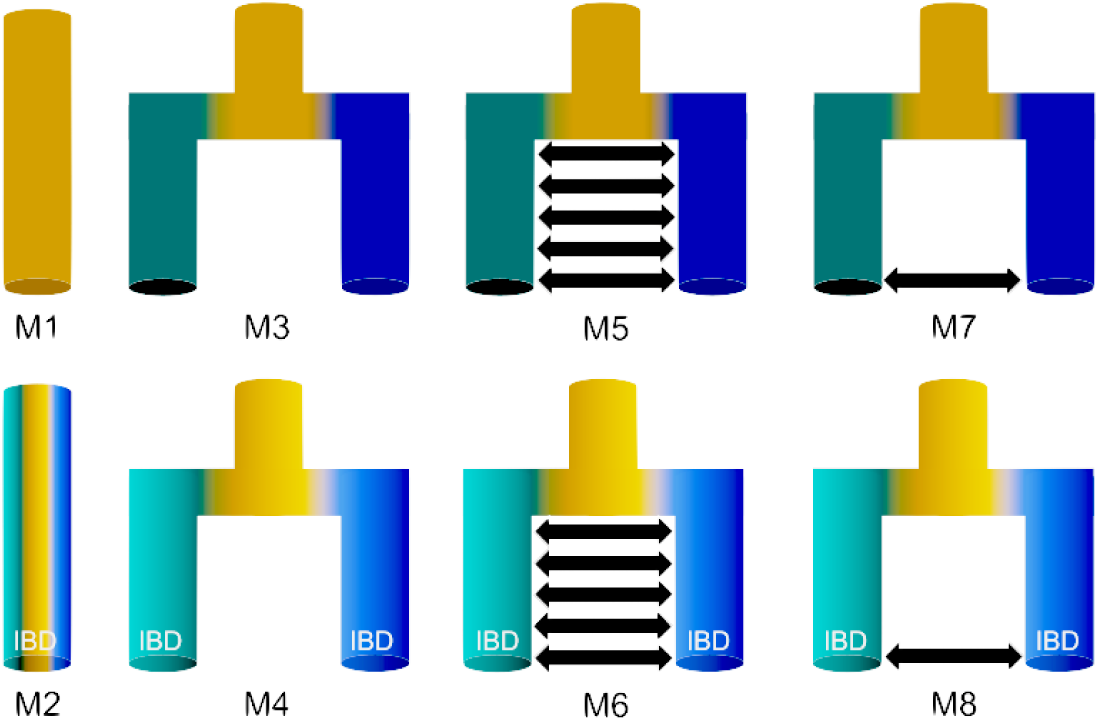
Visualization of the eight models simulated. Colorful tubes indicate populations. Arrows indicate gene flow. Color gradients within tubes indicate the presence of IBD, while solid colored tubes indicate no IBD. M1: Single population without IBD. M2: Single population with IBD. M3: Isolation without IBD. M4: Isolation with IBD. M5: isolation-with-migration without IBD. M6: isolation-with-migration with IBD. M7: Secondary contact without IBD. M8: Secondary contact with IBD.

Second, we simulated the presence of IBD to test whether it would impede our ability to determine phylogeographic structure. When IBD was implemented in a model, simulated individuals could only mate and produce offspring with their nearest neighbors. In isolation-with-migration models, gene flow could only occur between individuals close to the contact zone. In contrast, if IBD was not implemented, individuals could choose any other member of the population to produce offspring with, irrespective of geographic distance. These models were spatially-explicit with respect to both the environment that individuals were simulated on and mate choice and production of offspring (see Appendix 1).

We simulated four different lengths of time: 6,000, 21,000, 120,000, and 1,000,000 generations. These approximately correspond to the mid-Holocene, last interglacial optimum, last glacial maximum, and mid-Pleistocene, assuming a generation time of one year. We chose to use different time frames to 1) test if we could tell apart the different models when divergence times were short, 2) investigate divergence times that could have taken place during major environmental shifts in the CFB region, and reflect variation in the divergence dates found in our empirical analysis (see Results). This gave 32 simulation regimes in total, which were all combinations of the four demographic models, presence/absence of IBD, and four time frames.

We selected parameter values for the models based on empirical estimates and suggested practices (Table 1; see Appendix 1). We used ecological niche models (ENMs) to provide the landscape under which all demographic models were simulated. This is coarse-scale resolution, solely to generate a scenario where the area between the deserts is not as suitable as the area within the deserts. To do this we used an ENM of Bell’s Vireo (*Vireo bellii*), a species that is dependent on desert habitats, with known population structure and no evidence of dispersal across the CFB (Klicka, Kus, and Burns, 2016; see Appendix 1). From the simulated genetic data, we calculated 11 summary statistics using the R package ‘popgenome’ (Pfeifer, Wittelsbürger, Ramos-Onsins, and Lercher, 2014; Table S1.2). These include statistics known to be correlated with particular aspects of demographic histories (e.g., Tajima’s D, Tajima, 1989; haplotype diversity, Stumpf, 2004; see Appendix 1).

**Table 1:**
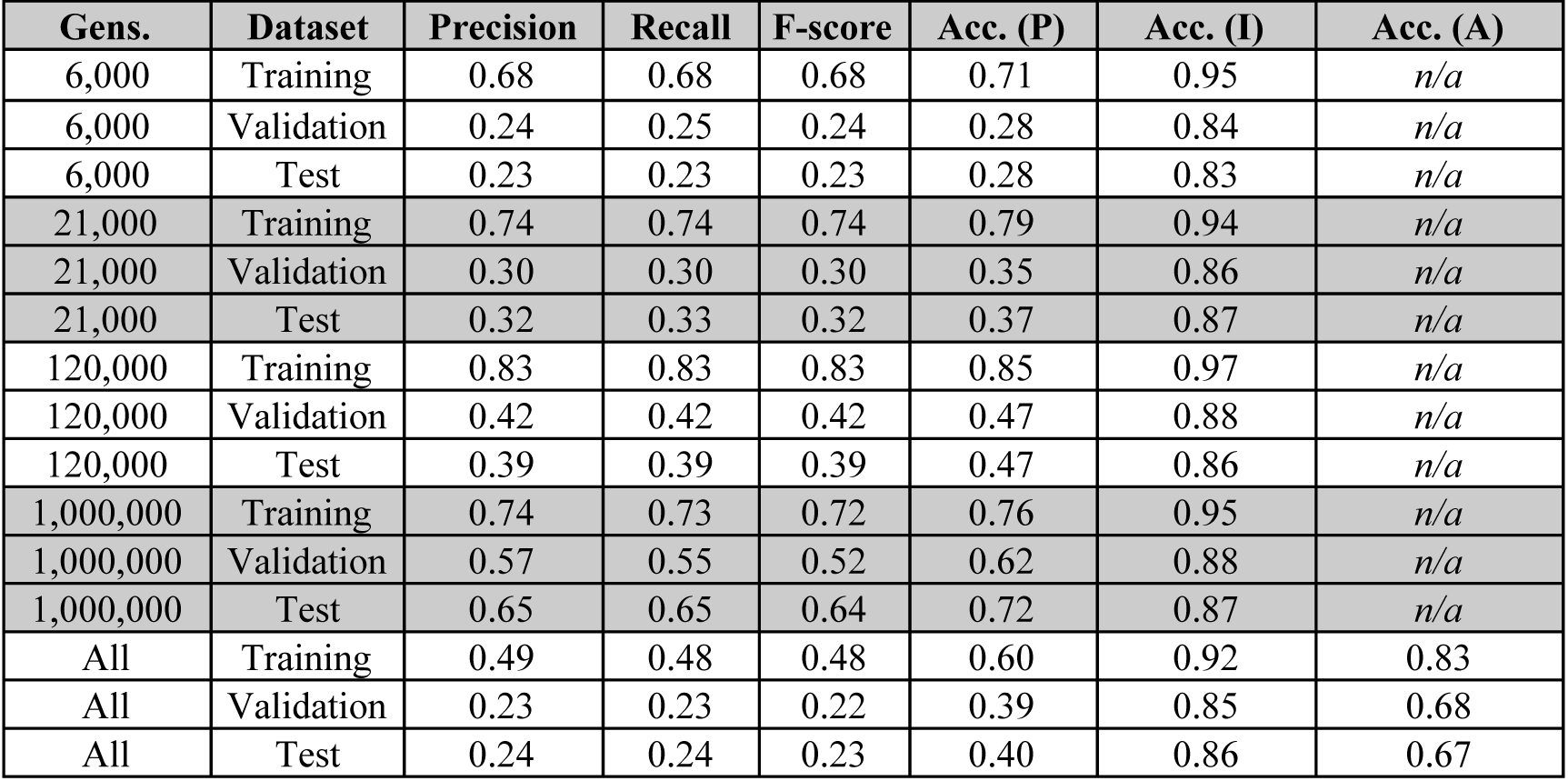
Model performance. Accuracies are calculated for phylogeographic structure alone (P), IBD alone (I), and ages alone (A) when applicable. “Gens.” =generations. “Acc.” =accuracy.

| **Gens.** | **Dataset** | **Precision** | **Recall** | **F-score** | **Acc. (P)** | **Acc. (I)** | **Acc. (A)** |
| --- | --- | --- | --- | --- | --- | --- | --- |
| 6,000 | Training | 0.68 | 0.68 | 0.68 | 0.71 | 0.95 | *n/a* |
| 6,000 | Validation | 0.24 | 0.25 | 0.24 | 0.28 | 0.84 | *n/a* |
| 6,000 | Test | 0.23 | 0.23 | 0.23 | 0.28 | 0.83 | *n/a* |
| 21,000 | Training | 0.74 | 0.74 | 0.74 | 0.79 | 0.94 | *n/a* |
| 21,000 | Validation | 0.30 | 0.30 | 0.30 | 0.35 | 0.86 | *n/a* |
| 21,000 | Test | 0.32 | 0.33 | 0.32 | 0.37 | 0.87 | *n/a* |
| 120,000 | Training | 0.83 | 0.83 | 0.83 | 0.85 | 0.97 | *n/a* |
| 120,000 | Validation | 0.42 | 0.42 | 0.42 | 0.47 | 0.88 | *n/a* |
| 120,000 | Test | 0.39 | 0.39 | 0.39 | 0.47 | 0.86 | *n/a* |
| 1,000,000 | Training | 0.74 | 0.73 | 0.72 | 0.76 | 0.95 | *n/a* |
| 1,000,000 | Validation | 0.57 | 0.55 | 0.52 | 0.62 | 0.88 | *n/a* |
| 1,000,000 | Test | 0.65 | 0.65 | 0.64 | 0.72 | 0.87 | *n/a* |
| All | Training | 0.49 | 0.48 | 0.48 | 0.60 | 0.92 | 0.83 |
| All | Validation | 0.23 | 0.23 | 0.22 | 0.39 | 0.85 | 0.68 |
| All | Test | 0.24 | 0.24 | 0.23 | 0.40 | 0.86 | 0.67 |

### Machine learning pipeline

We performed model selection across demographic regimes by building a neural network using ‘scikit-learn’, a Python module specifically for machine learning (Pedregosa et al., 2011). Model selection was performed so that we could tell whether the demographic model generating the data could be accurately detected. We analyzed each of the time periods together, as well as separately. The inputs were the 11 summary statistics outlined above (see Appendix 1). The network outputs a 3×1 matrix where each value in the matrix was the predicted population structure, predicted IBD value, and predicted divergence time. To evaluate the model performance we calculated Accuracy, Precision, Recall, and the F-score. Accuracy is the percentage of correctly identified positives. Precision is the number of true positives over the number of positives identified (i.e., true and false positives). Recall is the number of true positives over the number of actual positives (i.e., true positives and false negatives). The F-score is the weighted average of Precision and Recall.

## Results

### Biota around CFB shows variability in evolutionary histories

We found 99 published studies comprising 68 taxa (species complexes of 1–6 species) in 39 families and 19 orders (Asterales, Caryophyllales, Cucurbitales, Pinales, Araneae, Coleoptera, Hymenoptera, Orthoptera, Scorpiones, Anura, Squamata, Testudines, Galliformes, Passeriformes, Piciformes, Artiodactyla, Carnivora, Chiroptera, and Rodentia; Table S1.3; see Appendix 2 fordata sources). Phylogeographic inference was performed using a range of data including mitochondrial, chloroplast, and nuclear loci, microsatellites, allozymes, fragment length polymorphisms, and next generation sequencing (e.g., RADseq, UCEs). Some taxa had descriptions of variation across space that were based on phenotypic assessments, but all but one species had genetic data to reinforce those inferences. Seventeen taxa were supported using next-generation sequencing data.

The majority of the taxa (∼62%) showed evidence for phylogeographic structure across the CFB (Figure S1.1). Of the 68 taxa examined, 41 showed structure (with 38/41 monophyletic), 13 showed no structure, and the remainder had unclear results where it was ambiguous whether there was structure. Divergence times were estimated in 27/41 taxa and ranged from the Miocene to the Pleistocene with no clear temporal congruence across species. Of these, 12/27 were in the Pleistocene alone, with an additional three overlapping the Pleistocene and other epochs. The remaining 18 were not explicitly dated, but for five of those taxa Pleistocene glacial cycles are cited as being a major driver of divergence.

Just under half of the taxa with phylogeographic structure had an estimate of gene flow (Table S1.3, Figure S1.1). Of the 32 taxa, five had no gene flow across the CFB, 13 had gene flow, and the rest had ambiguous results. For the 17 species that were structured and had gene flow estimated, these numbers were three with no gene flow, seven with gene flow, and seven with ambiguous results. Allopatry was the primary mode of speciation proposed, with isolation-by-environment secondary. Some of the studies that found support for gene flow (and thereby isolation-with-migration or secondary contact) concluded that pure allopatric speciation was taking place. Of the 55/68 taxa that had clear or ambiguous splits across the CFB, allopatric speciation was the mode of speciation proposed for 19 taxa, with an additional five identifying allopatry with another mode of speciation, including hybrid speciation, polyploidy, and sexual selection (Table S1.3). Another 12/54 declared isolation-by-environment alone (often with the influence of IBD) as the main driver; all these taxa had support from next-generation-sequencing data and were from the same study (Myers et al., 2019). The remaining 19 taxa did not have inferred modes of speciation. There appears to be a temporal bias in the interpretation of these results: only recent papers (2004–2019, median 2017) suggested isolation-by-environment as the main driver, while older papers overwhelmingly suggested allopatry either alone (1986–2018, median 2005) or with another mechanism (1996–2014, median 2005).

The location and width of the contact zone varied between species and also varied with respect to the divergence times and locomotor types (Figure 3; Figure S1.2, Figure S1.3). The total extent of the barrier ranged from 118.3–99.5°W longitude, or 18.8° longitude width. Of the 36 species for which we could estimate where the barrier was, 24/36 overlapped 108.6°W, the maximum number of species to overlap. The zone of overlap for over 50% of the taxa (18 or more species) ranged from 109.3–105.6°W. The contact zone widths for each taxon ranged from 0.3–11.2°W.

**Figure 3:**
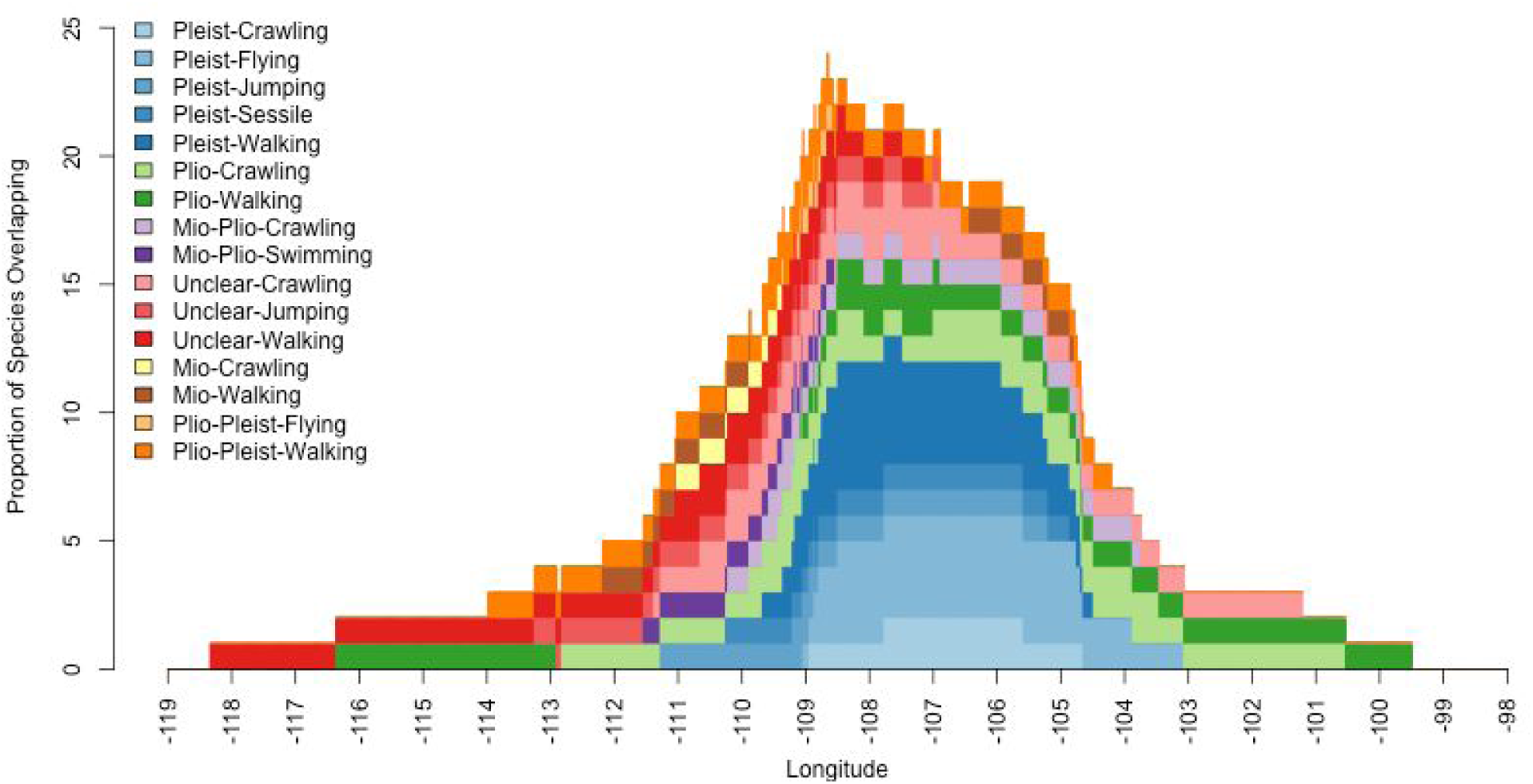
The width of the CFB contact zone for taxa varies by the timing of divergence and the type of locomotion. Colors correspond to unique combinations of divergence and locomotion. Hues represent divergence times (Pleistocene, Pliocene, Miocene, Unclear, etc.) and shades represent locomotion types (Crawling, Flying, Walking, etc.). X-axis shows the longitudinal range, Y-axis shows the number of species that overlap a given longitude. See also Figures S1.2 and S1.3.

The overall width of the contact zone at the barrier varies with respect to divergence time (Figure S1.2). Width peaks for taxa who diverged in the Pliocene (16.9°), decreasing as divergence dates get both older (7.4–7.7°) and younger (8.2–9.3°). The contact zone width also varies with respect to locomotion type (Figure S1.3; see Appendix 1). Walking taxa span the entire range of 18.8°. For the other locomotion types, the largest to smallest width goes crawling, jumping, flying, sessile, and swimming taxa (12.7–3.0°). In addition, locomotion type and structure co-vary (Figure 4). Flying species have the lowest percent of taxa with structure (8/17, ∼47%). All swimming and jumping taxa (*N*=1, *N*=3) have structure. Of the rest, walking species have the highest percentage of taxa with structure (19/27, ∼70%) followed by crawling (10/16, ∼63%) and sessile species (2/4, 50%).

**Figure 4:**
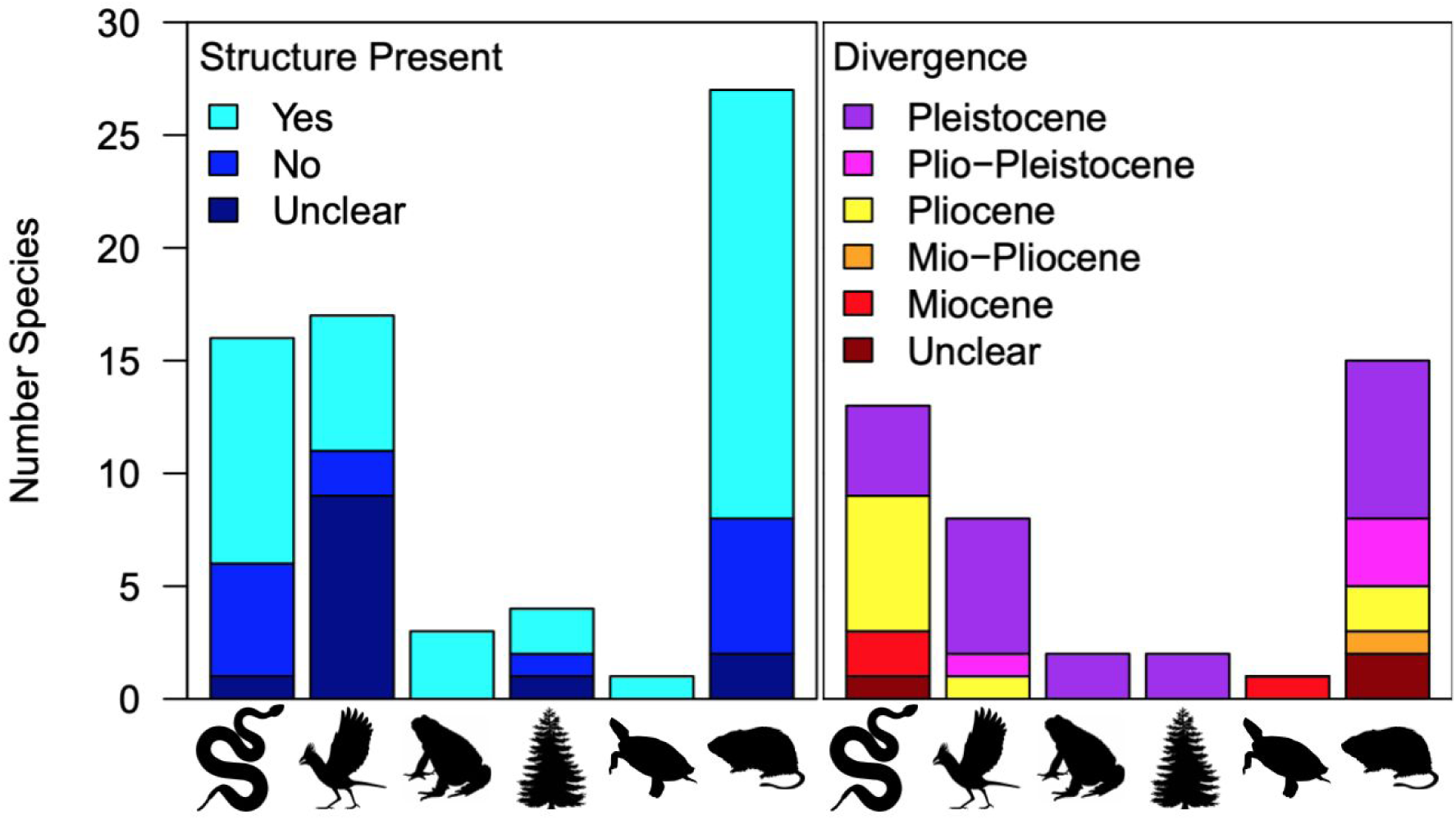
The presence of phylogeographic structure and divergence time across the Cochise Filter Barrier vary based on the locomotion type. X-axis: locomotion type (from left: crawling, flying, jumping, sessile, swimming, walking). Y-axis: number of species per category. Colors describe the presence/absence of phylogeographic structure (left), or the divergence time (right).

The influence of thermoregulation on divergence appears to be less clear than the influence of locomotion (see Appendix 1). A higher proportion of ectotherms are found to have structure (27/40, ∼68%) than endotherms (14/28, 50%), but the difference is not significant (χ^2^=2.11, p=0.15). Ectotherms may also have older divergences: all of the taxa whose divergences overlap the Miocene are ectothermic (Figure S1.4) and of the remaining, proportionately fewer endotherms diverge in the Pliocene (2/9, ∼22%) than the Pleistocene (14/27, ∼52%). Both thermoregulation and locomotion are significant predictors of divergence date (thermoregulation R^2^=0.14, p=0.0078, locomotion R^2^=0.25, p=0.025), though not when they are both in the same model (R^2^=0.31, thermoregulation p=0.060, locomotion p=0.088). They are still significant when accounting for elevational preference, taxonomy, or both simultaneously (range R^2^=0.14–0.34, thermoregulation p=0.0054–0.014, locomotion p=0.013–0.047). Of the taxa examined, 34 were found in lowlands, 10 were montane, and the remainder were found in both; however, elevational preference is not a significant predictor of divergence time (with and without taxonomy p=0.57).

### Accuracy of machine learning classification varies with age and phylogeographic history

After calculating summary statistics, removing duplicate simulations that produced the exact same statistics, and eliminating collinear variables, we had 26,131 unique sets of summary statistics. The number of summary statistic sets per model varied based on the number of generations each model ran for (6,000 generations: 834–1,099 sets per model; 21,000: 1,011–1,200; 120,000: 1,100–1,100, 1,000,000: 80–100). Generally, younger runs were more likely to produce duplicate results.

The neural network varied in accuracy (Table 1, Table S1.4), but was consistently good at classifying whether models had IBD, less good (but consistent) at classifying the number of generations after divergence, and inconsistent at classifying the true demographic model. When the network was asked to classify if IBD was present, it did so with high accuracy regardless of how many generations after divergence or how many summary statistics were used (92–100% for training, 81–89% for validation/testing), and overfitting appeared to be slight (Table 1, Table S1.4; Figure S1.5). Similarly, when the network classified the number of generations since divergence, it did so with 83–85% accuracy in training, depending on how many summary statistics were used, and 66–68% in validation/testing. When models were misclassified, it was always to a similar number of generations (e.g., 21,000 generation runs were confused with 6,000 and 120,000 generation runs, but never 1,000,000 generation runs; Figure S1.5). The differences in training and validation/testing suggest slight overfitting. Finally, neural network accuracy for classifying phylogeographic models was much more variable with a lot of overfitting (61–94% training, 28–62% validation/testing). Accuracy was positively associated with the number of generations after divergence models were run for, with those that ran for longer having higher ability to differentiate between models (Table 1, Table S1.4).

When examining performance across models (Figure 5), IBD had similar misclassification rates irrespective of the number of generations since divergence and phylogeographic structure simulated. The exception is for models run for 1,000,000 generations after divergence. Here, the single population with IBD models had high (worse than random) misclassification rates. Otherwise, models run for 6,000-120,000 generations tend to have higher misclassification rates, though the trend is reversed for secondary contact models, which have lower misclassification rates. Phylogeographic structure was also more highly misclassified in models with a short number of generations since divergence, irrespective of IBD. Multiple models are classified worse than randomly based on age: single population with IBD models (1,000,000 generations), isolation with IBD models (6,000 generations), and isolation-with-migration both with (21,000 generations) and without IBD (6,000 generations, 21,000 generations).

**Figure 5:**
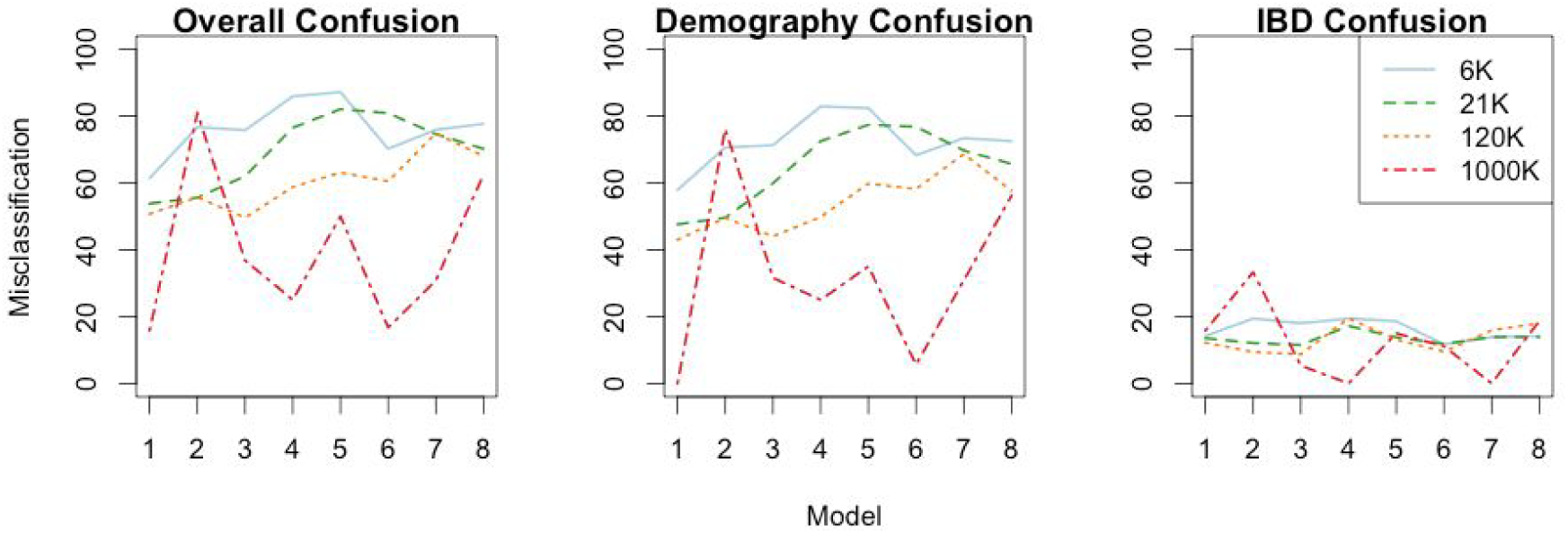
Neural networks classify models with older divergences much better than models with younger divergences. Misclassification rates are given for overall models, phylogeographic models only, and IBD models only. Model numbers on the x-axis correspond to models in Figure 2. Y-axis indicates misclassification rate, with higher values indicating poorer model performance. Lines of different colors and dash types indicate the generations simulated, from 6,000 (6K) to 1,000,000 (1000K) generations.

We examined which misclassifications were most common with respect to the number of generations modeled since divergence (Figure 6, Figure S1.6). From a phylogeographic structure perspective (irrespective of IBD), across all ages of divergence, single population models were likely to be misclassified as isolation-with-migration models and vice versa. Likewise, isolation models were likely to be misclassified as secondary contact models and vice versa. In models that were run for 1,000,000 generations since divergence, misclassifications between isolation/secondary contact and isolation-with-migration/single population were the only misclassifications that occured. However, as the number of generations since divergence decreased, more kinds of misclassifications arose. In models run for 120,000 generations after divergence, isolation-with-migration and secondary contact were misclassified, as were single population and secondary contact. All models were confusable in younger divergences, with confusion greatest in models run 6,000 years after divergence. This reveals that even in lineages with old divergences, single population and isolation-with-migration models are hard to tell apart, as are isolation and secondary contact models. However, the younger a divergence is, and the less time for genetic differences to accumulate, the easier it is to confuse all types of phylogeographic models. When considering misclassifications simultaneously between IBD and phylogeographic structure models, in older models we generally find that when models are misclassified between having IBD and not having IBD, it tends to be within the same model type (i.e., isolation-with-migration with IBD to isolation-with-migration without IBD) or between those models that were already identified as similar when not considering IBD (i.e., secondary contact with IBD to isolation without IBD). However, as models become younger, misclassification becomes more severe.

**Figure 6:**
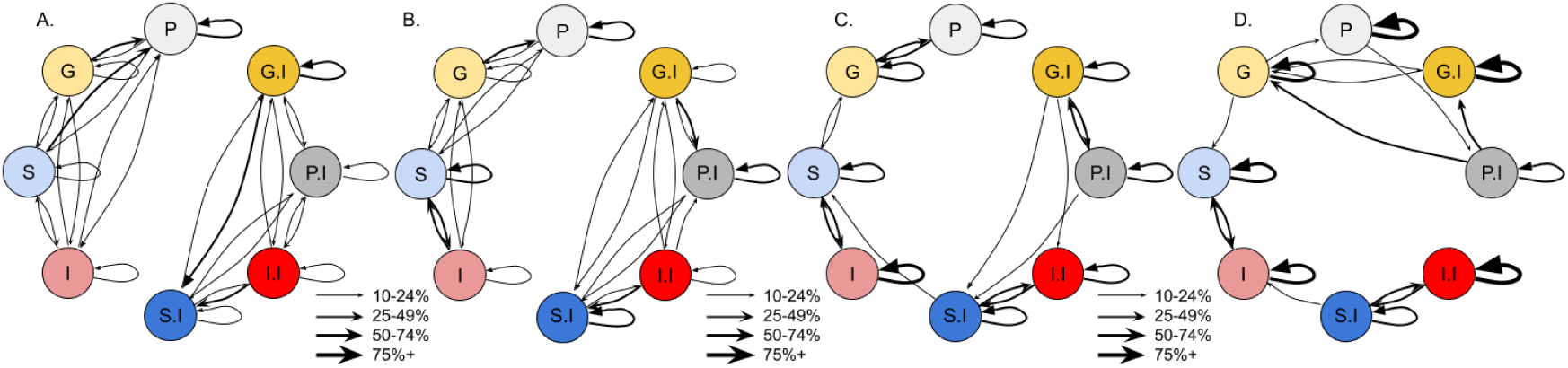
Overall model classifications are easily confused at young ages but improve with older ages. Arrows begin at the true model and end at the assigned model. Line thickness of arrows indicates the percentage of time those assignments are made. Assignments under 10% are omitted for clarity. Demographics include single panmictic populations (“P”, grey), isolation-with-migration or gene flow (“G”, yellow), secondary contact (“S”, blue), and isolation (“I”, red). Suffix of “.I” after demography indicates that IBD is present. A) 6,000 generations. B) 21,000 generations. C) 120,000 generations. D) 1,000,000 generations. Models with IBD are on the right, models without IBD are on the left.

## Discussion

Moving beyond descriptive narratives of landscape change generating biodiversity and understanding how taxa interact with barriers represent major challenges of biogeographic research. We used the Sonoarn and Chihuahuan Deserts as an exemplar system for characterizing genetic differentiation among two regions, quantifying the temporal and spatial dynamics in phylogeographic structuring associated with a filter barrier. The range in divergence times and phylogeographic break locations in taxa distributed across the deserts reflects the semipermeability of the CFB. The variability of molecular data used in this synthesis was not suitable for a formal test of simultaneous divergence (e.g., Hickerson et al., 2006), but the wide disparity of divergence times suggests taxa have been diverging across the CFB for millions of years. Locomotive and thermoregulatory traits help to explain this variation, suggesting that organisms that are more vagile and less dependent on external temperatures (i.e., endotherms) may be better at evading the impacts of the barrier, and selective pressures both on movement and on environmental tolerances might determine which taxa diverge. Divergence in allopatry was typically invoked as the main mode of differentiation across the CFB, but we found evidence of all other processes (i.e., ecological divergence, polyploidy, hybrid speciation, and sexual selection). Fully understanding which filters make up the CFB necessitates a deeper understanding of the mechanisms causing differentiation in the region. Using simulations and model selection approaches can illuminate those mechanisms via rigorous hypothesis testing and replication. These methods offer promise by more accurately identifying modes of differentiation, but the modeled scenarios can be difficult to distinguish. Nevertheless, simulation methods contextualize empirical results and show the utility of genomic-scale data in uncovering phylogeographic histories, allowing us to understand how organisms might be expected to diversify under different known conditions.

### The community around barriers are influenced by a multitude of forces

Biogeographic barriers are typically viewed as abrupt transitions between two biotas, but our analysis showed the conditions in which this view does not hold. The entities identified as biogeographic barriers may represent a combination of forces that generated the contemporary distributions of taxa, which have a wide longitudinal range of phylogeographic breaks across the CFB after accounting for uncertainty. Most species have ranges that overlap ∼109–106, a shift eastward from the ∼112–108 that was conventional in the literature (though very few species do not overlap the original distribution). First, barriers can be ephemeral over time, for example, due to river capture events (Tagliacollo, Roxo, Duke-Sylvester, Oliveira, and Albert, 2015), sea-level changes (Elias, Short, Nelson, and Birks, 1996), or major climatic shifts (possibly including the CFB; Holmgren et al., 2007). Additionally, organisms have the capacity to disperse, which can retain or reintroduce gene flow and connectivity across barriers (Brandley, Guiher, Pyron, Winne, and Burbrink, 2010). Population genetics theory also predicts that even in the absence of a barrier, phylogeographic structure can arise due to simple density troughs, where abundances are low and dispersal out of these troughs fails (Barton and Hewitt, 1981; Barrowclough et al., 2005). Finally, incomplete sampling may be a culprit for the observed differences between taxa, as contact zone widths cannot be estimated without examining specimens. For the CFB, numerous studies lack individuals from the transition zone. Fortunately, targeted sampling of this region will resolve this discrepancy, especially with the addition of genome-level genetic sampling. In our analyses here, we used all individuals to calculate summary statistics, but future work could subset individuals and create incomplete sampling to test for bias.

### Locomotion and thermoregulation as filters

Our results show the expected relationship between locomotion type and how organisms respond to the barrier. We found that flying organisms are less likely to have diverged across the barrier. Flying organisms can have higher dispersal ability than non-flying ones, allowing them to potentially bypass any filtering mechanisms of a biogeographic filter barrier, and previous work has shown that flying organisms tend to show less genetic differentiation than others (Medina, Cooke, and Ord, 2018). The contact zone at the barrier also has a smaller estimated width for flying taxa than those with other forms of locomotion. Under an assumption where flying confers higher dispersal, it is counterintuitive that they should show a narrow range for the CFB, unless dispersal is unable to counteract the strength of the filters operating, but the degree of environmental changes may be relatively stronger than the effects of organismal traits. Flying organisms have a higher proportion of Pleistocene divergences relative to other epochs even after accounting for elevation, which has been found across other biogeographic barriers (e.g., Bacon et al., 2015). It is unclear whether the Pleistocene climatic shifts themselves caused diversification or if some other process produced a suite of divergences that happen to date to the Pleistocene for other reasons. Alternatively, flying species may have more suitable habitat available to them across the barrier relative to other taxa, which would weaken the impact of the CFB. This would explain both the shallow divergences and the narrow width. In addition, some taxa are clearly undersampled (e.g., sessile plants) or less abundant in desert habitats (e.g., swimming fish), though for other taxa it is less clear. Increasing sampling of organisms with these forms of dispersal would clarify the causes of these trends.

Ectothermic taxa appear more likely to diversify across the CFB than endothermic taxa, though the patterns are not significant. As ectothermic organisms are more reliant on external temperature regulation than endotherms (particularly microhabitat selection; Boyko, 2014), it is feasible that any environmental changes should disproportionately affect them. Under such an assumption, the climatic dynamism of the Pleistocene would have been a stronger filter on ectotherms, though it would not explain the ectothermic taxa that diverged in the Pliocene. Alternatively, local adaptation could be a stronger process in ectotherms relative to endotherms, which would explain divergence under an ecological-speciation hypothesis (Nosil, 2012).

The traits we chose to use in this study were simple to quantify across all of these taxa. Many more traits may be involved in mediating which species diverge across this and other barriers. For example, habitat preferences could be driving the pattern in addition to or instead of the ones we have identified here. The relative width of the contact zone for a taxon could be changed as the traits interact with the properties of the barrier, for example, whether it allows frequent or infrequent dispersal (see Pyron and Burbrink, 2010), is long-lived or ephemeral, and/or relocates through time. Further, both of these traits are presumably the result of selection, whereas more stochastic forces could be at play. Overall, phenotypic traits could be important for determining why multiple co-distributed taxa are concordant or discordant with respect to their phylogeographic histories (Zamudio, Bell, and Mason, 2016).

One major limitation of this study is that it does not assess any taxa that completely fail to get through the barrier, i.e., taxa that are restricted to one side. There are numerous examples of single-desert endemics in the region, one of the most famous being that of the saguaro cactus (*Carnegiea gigantea*) of the Sonoran desert. Understanding which organisms fall into that category would greatly improve knowledge of what factors make the CFB a potent barrier. A difficulty in such an assessment would be identifying which organisms have successfully crossed but one population was extirpated by chance vs. organisms that never crossed successfully. Still, the taxa that are unable to get past the barrier would give insight into exactly what filters are operating through analyses of their traits and the properties of the barrier.

### Genome-level data can clarify phylogeographic patterns

Ultimately, even in aggregate, the studies performed on the CFB do not provide enough information to explain the mechanisms of diversification in the region. Using machine learning methods on large simulated datasets, combined with empirical and theoretical studies, allows for testing between phylogeographic models with numerous parameters of interest; for example, the divergence mechanisms in Australian honeyeaters are ambiguous in empirical data but may be resolvable using a simulated approach (Toon, Hughes, Joseph, 2020). Our simulations here are only a subset of model parameter space and larger genomic regions, population growth, and selection could be considered (Carstens et al., 2013; Ewing and Jensen, 2016). Likewise, we do not account for changes in parameter space over time (e.g., population size, mutation rate, environmental factors). Demographic expansions and contractions in effective population size cause changes in the impact of genetic drift, and there are numerous examples of phylogeographic studies that find these changes in concordance with genetic structure (e.g., Smith et al., 2011; Charruau et al., 2012). Changes in habitat suitability over time can lead to similar changes; fortunately, spatially explicit methods can incorporate these changes via the addition of paleoclimate models, some of which go back to the Pliocene (e.g., PRISM4; Dowsett et al., 2016).

The machine learning method we used here, neural networks, allow for direct estimation of the model power and model identifiability in test data, which can be leveraged when analyzing both simulated and empirical datasets. Validation schemes indicate how well a model performs on known models, giving estimates of error and describing which models have problems with identifiability. Here, the identifiability of our models was high when classifying IBD and generation times. However, it is notable that at very young divergence times, our method still struggled to tell apart different models, which may imply that differences between isolation and secondary-contact models are inscrutable without longer divergence histories. The difficulty in identifying models is well known when analyzing certain data types, the SFS in particular: population size is hard to estimate, especially when it has been dynamic over time (Lapierre, Lambert, and Achaz, 2017; Terhorst and Song, 2015), and the presence of ancient population structure can erroneously appear as though gene flow has occurred (Eriksson and Manica, 2014). If these tenets also hold for additional summary statistics, they must be accounted for in future work. Nevertheless, this all serves to reinforce that biologically similar models tend to be confused with one another (e.g., Roux et al., 2016). Further work will require more rigorous testing as to the impact of model selection on different biological scenarios.

In conclusion, we synthesized phylogeographic studies across an exemplar barrier, examining how biogeographic filters affect the community. Among all species, gene flow, phylogeographic structuring, and timing of divergence all varied substantially. The location of the barrier and width of the contact zone are related to locomotion and thermoregulation, irrespective of phylogeny and elevational preferences. Simulations and machine learning allowed us to quantify spatiotemporal evolutionary histories, finding numerous aspects of demography including gene flow and isolation-by-distance are major confounding variables. Overall, barriers interact with traits of species to cause heterogeneous differentiation, but identifying the exact causes remains challenging.

## Acknowledgements

We are grateful to all the authors whose work supported our synthesis. KLP was supported by Frank M. Chapman, Sydney Anderson, and Linda H. Gormezano funds, AOS, SSB, and RGGS. Computational resources provided by the Open Science Grid (Pordes et al., 2007, Sfiligoi et al., 2009; NSF award 1148698 and the U.S. DoE’s Office of Science). EAM was supported through the Peter Buck and Rathbone Bacon Fellowship from the National Museum of Natural History. We thank F. Burbrink for the photographs. Feedback comes from M. Blair, L. Alter, C. Raxworthy, J. Padial, E. Rodriguez, L. Musher, L. Moreira, J. Merwin, G. Thom, G. Seeholzer, V. Chua, J. Denton, D. Kelly, I. Overcast, A. Xue, M. Ingala, M. Hickerson, M. Gehara, and B. Haller.

## Appendix 1: Supplementary Analyses and Results

We classified taxa into categories based on published elevational ranges: desert lowland, highland montane, or both. We then performed GLMM to predict divergence times based on combinations of 1-3 organismal traits (locomotive, thermoregulatory, elevational), and one of six classification groups (i.e., Genus through Kingdom) as a random effect. Thus we ran 49 models representing these combinations. Elevation is never a significant predictor of divergence time in any model, before or after accounting for taxonomy. Before accounting for taxonomy, thermoregulation and locomotion are significant irrespective of elevation. However, models accounting for both thermoregulation and locomotion simultaneously are never significant. The models that are significant before accounting for taxonomy remain significant when Genus, Family, Phylum, or Kingdom ranks are used as the random effect. However, they are not significant when using Order or Class as the random effect. Our interpretation of these results is that there are too many Genera and Families relative to data points, and too few Phyla and Kingdoms, to act as effective random effects. Therefore, this shows evidence that thermoregulation and locomotion are not significant after accounting for taxonomic levels.

### Niche Modeling

We obtained occurrence data for *Vireo bellii* by downloading all eBird data in May 2017 (Sullivan et al., 2009). We then supplemented this data with data from GBIF, iNaturalist, and VertNet (gbif.org, inaturalist.org, vertnet.org). This was done using custom R scripts that made use of the following packages: ‘raster’ (Hijmans 2017), ‘MASS’ (Venables and Ripley 2002), ‘spocc’ (Chamberlain 2017), ‘rgeos’ (Bivand and Rundel 2017), ‘dplyr’ (Wickham et al., 2017), ‘sp’ (Pebesma and Bivand 2005; Bivand et al., 2013), and ‘dismo’ (Hijmans et al., 2017). We removed duplicate localities. Outliers were detected and removed using a probability density function. After this, we thinned the data using ‘spthin’ (Aiello-Lammens et al., 2014) to account for spatial autocorrelation in the data. This left us with 1,026 occurrence points for the species.

Climate data used was the WorldClim database (Hijmans et al., 2005). We used all 19 variables, but we constrained the climate data to fall within longitude −115.70 to −101.26 and latitude 27.00 to 36.47. We then took the climate data and the occurrence data and built an ecological niche model using MaxEnt (Phillips et al., 2006) with ‘ENMeval’ as a wrapper function for model selection (Muscarella et al., 2014). ‘ENMeval’ is an R package that optimizes MaxEnt models based on different sets of feature classes and regularization values. In particular, we selected between the linear and linear+quadratic feature classes, as well as regularization values of 0.5, 1, 1.5, 2, 2.5, 3, 3.5, and 4. Models were run for each combination of feature classes and regularization values, and the model with the lowest occurrence point omission rate and highest area under the curve was chosen. The best model used linear+quadratic feature classes and a regularization value of 0.5, with an AUC of 0.83 and an omission rate of 14.2%. The second best model also used linear+quadratic features, but had a regularization value of 1, an AUC of 0.82, and an omission rate of 22.8%. After the final model was generated and projected back onto the climate data, we converted the suitability values from 0-1 for SLiM to use and downprojected to an ascii of 57 by 87 grid cells.

### Slim Simulation Details

For each of the 32 model sets, we ran 1000-1200 simulations each in SLiM 3 with the exception of the eight models run for 1,000,000 generations. These longer models were computationally intense, and so we only ran them 100 times each, to determine whether the patterns we found for 6,000-120,000 generations would continue to hold for longer time periods. Scripts are available at github.com/kaiyaprovost.

We used a mutation rate of 2.21 × 10^−9^ substitutions per site per year (Nam et al., 2010), the suggested default recombination rate in SLiM 3.1, and empirical estimates of ancestral population sizes for a bird with genetic structure across the CFB (Provost et al., 2018). We chose a large gene flow value of 10% to amplify the effects of introgression within applicable models. We simulated a chromosome of 100,000 base pairs. For models with a phylogeographic split, we divided individuals into two equally sized populations. The spatial boundaries of these populations were constrained by longitude to simulate a barrier; individuals in population 1 were only able to occupy the western 50% of habitat, and individuals in population 2 were only able to occupy the eastern 50% (see below).

The niche model was constrained to be between longitude −115.70 to −101.26, and latitude 27.00 to 36.47, narrowing in on the transition zone between the deserts. Fitness across the landscape was calculated by taking the cube root of the cell value (which ranged from 0–1) and multiplying it by the relative fitness of the individual; these fitness values are required in SLiM to create new individuals each generation and cause the algorithm to preferentially generate offspring in regions of high ecological niche suitability.

We reduced the computational load of our models by scaling our parameters by a factor (lambda) of 0.02, and by implementing tree sequence recording (Haller et al., 2019). Scaling factors reduce the number of generations and number of individuals to simulate but keep the relative mutation rate and recombination rate constant. We chose 0.02 as it improved our performance without impacting the distribution of summary statistics in preliminary runs (Table S1.1). Tree sequence recording tracks the true ancestry of every position in the simulated chromosome across individuals rather than explicitly simulating mutations (Haller et al., 2019). Individuals who do not have any descendents are removed. We modeled neutral variation from the tree sequences using our scaled mutation rate, then converted the tree sequences to VCF format (Danecek et al., 2011) and MS format (Stevison 2014). MS format summarizes the number of segregating sites, the positions with polymorphic sites, and the haplotypes of every individual where a zero indicates the ancestral state and a 1 indicates the derived state. When converting to MS format, we randomly sampled 20 diploid individuals, 10 from the eastern side of the barrier and 10 from the western.

We generated 33 summary statistics originally (Table S1.2). Many of these summary statistics are highly correlated with each other. There is some evidence to suggest that neural networks can suffer from multicollinearity issues (Cheng et al., 2018). As such we ran the neural network with all summary statistics as well as with a reduced set that only contained statistics correlated with an absolute correlation coefficient less than 0.75. This gave us 11 summary statistics (Table S1.2). In some cases, simulations produced duplicate summary statistic information, in that independent runs of the simulation generated the exact same 33 summary statistics upon output, likely because the same random seed was generated. We removed these duplicates going forward to avoid overfitting.

### Machine Learning Details

Our neural network used the LBFGS (Byrd et al., 1995) algorithm as its solver and had an adaptive learning rate, where the rate of learning decreased as the iterations increased. The learning rate was initialized at 0.0001. There were 1,000 maximum iterations allowed. The hidden layer was constrained to be three layers each of 100 nodes, which was chosen after testing between it and eight other configurations (one layer of five, 25, and 100 nodes; two layers of five, 25, and 100 nodes; three layers of five, 25, and 100 nodes) to determine which performed best at classifying our validation data. The alpha parameter, which penalizes complexity, was tested across values 0, 0.01, 0.1, 1, and 10; to choose between these, we ran the neural network five times while varying the alpha value, and the network with the highest accuracy was chosen.

Running the neural networks with the 33 summary statistics rather than the 11 summary statistics marginally changed our models; they decreased model accuracy across all model runs, improved compute time, and resulted in less overfitting. As such we only present model results using 11 summary statistics unless otherwise specified.

Machine learning algorithms have broadly been shown to have high accuracy in classifying models. The suite tools developed to utilize these algorithms have been used broadly in biological contexts, for example to aid in sound identification (Juang and Chen 2007) and image processing (Wei et al., 2018) as well as for the classification purposes we use them for here (Sukumaran et al., 2016). Despite these strengths, it is not known how well the simulations classify actual genetic data, as that was outside the scope of this study. Further, regardless of how hypotheses are generated and tested, some biological scenarios will still be difficult to tell apart even with powerful and state-of-the-art simulation software and analytical techniques (see main text; Roux et al., 2016).

### Supplementary Figures and Tables

**Table S1.1:** Parameters used in the simulated demographic models. “Gens” =generations. Scaled with a lambda=0.02.

| Parameter | Original Value | Scaled Value |
| --- | --- | --- |
| Effective Population Size ($N_e$) | 400,000 | 8,000 |
| Mutation rate (per base per gen) | $2.21 \times 10^{-9}$ | $1.11 \times 10^{-7}$ |
| Recombination (per base per gen) | $1 \times 10^{-8}$ | $5 \times 10^{-7}$ |
| Number of Gens. | 6,000 | 120 |
|  | 21,000 | 420 |
|  | 120,000 | 2,400 |
|  | 1,000,000 | 20,000 |
| Gens. of Secondary Contact | 1,000 | 20 |

**Table S1.2:** Summary statistics used as inputs to our neural network. Asterisks next to variable names indicate that the variable was kept in the 11-statistic, non-collinear dataset. “P1” and “P2” stand for population 1 and population 2, respectively.

| Variable Name | Description |
| --- | --- |
| div_per_site* | Calculated nucleotide diversity per site |
| fu.li.d* | Fu and Li’s D |
| fu.li.d.1 | Above, but calculated assuming two populations |
| fu.li.f | Fu and Li’s F |
| fu.li.f.1 | Above, but calculated assuming two populations |
| hap.diversity.within* | Haplotype diversity within populations |
| kurt_haplotype.counts* | Kurtosis of all frequencies of haplotype counts |
| kurt_minor.allele.freqs | Kurtosis of all minor allele frequencies |
| mean_haplotype.counts* | Mean of all frequencies of haplotype counts |
| mean_minor.allele.freqs | Mean of all minor allele frequencies |
| mean_nuc.diversity.within | Mean of nucleotide diversity within populations |
| n.biallelic.sites | The number of biallelic sites |
| n.segregating.sites* | The number of segregating sites in P1 |
| n.segregating.sites.1 | Above, but for P2 |
| n.sites* | The number of sites total |
| pi | Nucleotide diversity |
| skew_haplotype.counts | Skew of all frequencies of haplotype counts |
| skew_minor.allele.freqs* | Skew of all minor allele frequencies |
| skew_nuc.diversity.within | Skew of nucleotide diversity within populations |
| tajima.d | Tajima’s D for P1 |
| tajima.d.1* | Above, but for P2 |
| theta_achaz.tajima | Achaz’s Tajima’s D accounting for sequencing error in P1 |
| theta_achaz.tajima.1* | Above, but for P2 |
| theta_achaz.watterson | Achaz’s Watterson’s Theta accounting for error in P1 |
| theta_achaz.watterson.1 | Above, but for P2 |
| theta_fu.li | Fu and Li’s Theta in P1 |
| theta_fu.li.1* | Above, but for P2 |
| theta_tajima | Tajima’s Theta |
| theta_watterson | Watterson’s Theta in P1 |
| theta_watterson.1 | Above, but for P2 |
| var_haplotype.counts | Variance of all frequencies of haplotype counts |
| var_minor.allele.freqs | Variance of all minor allele frequencies |
| var_nuc.diversity.within | Variance of nucleotide diversity within populations |

**Table S1.3:**
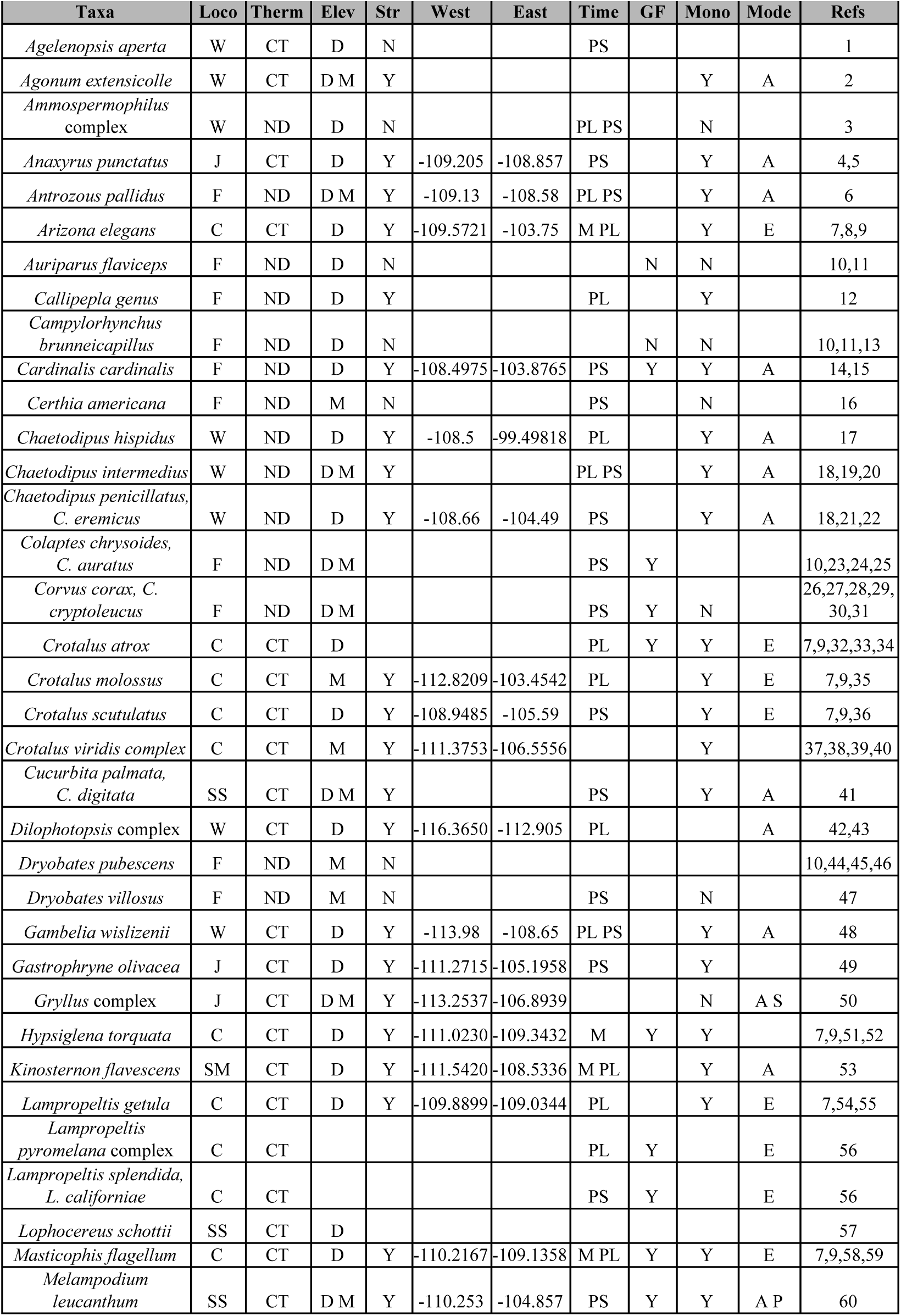
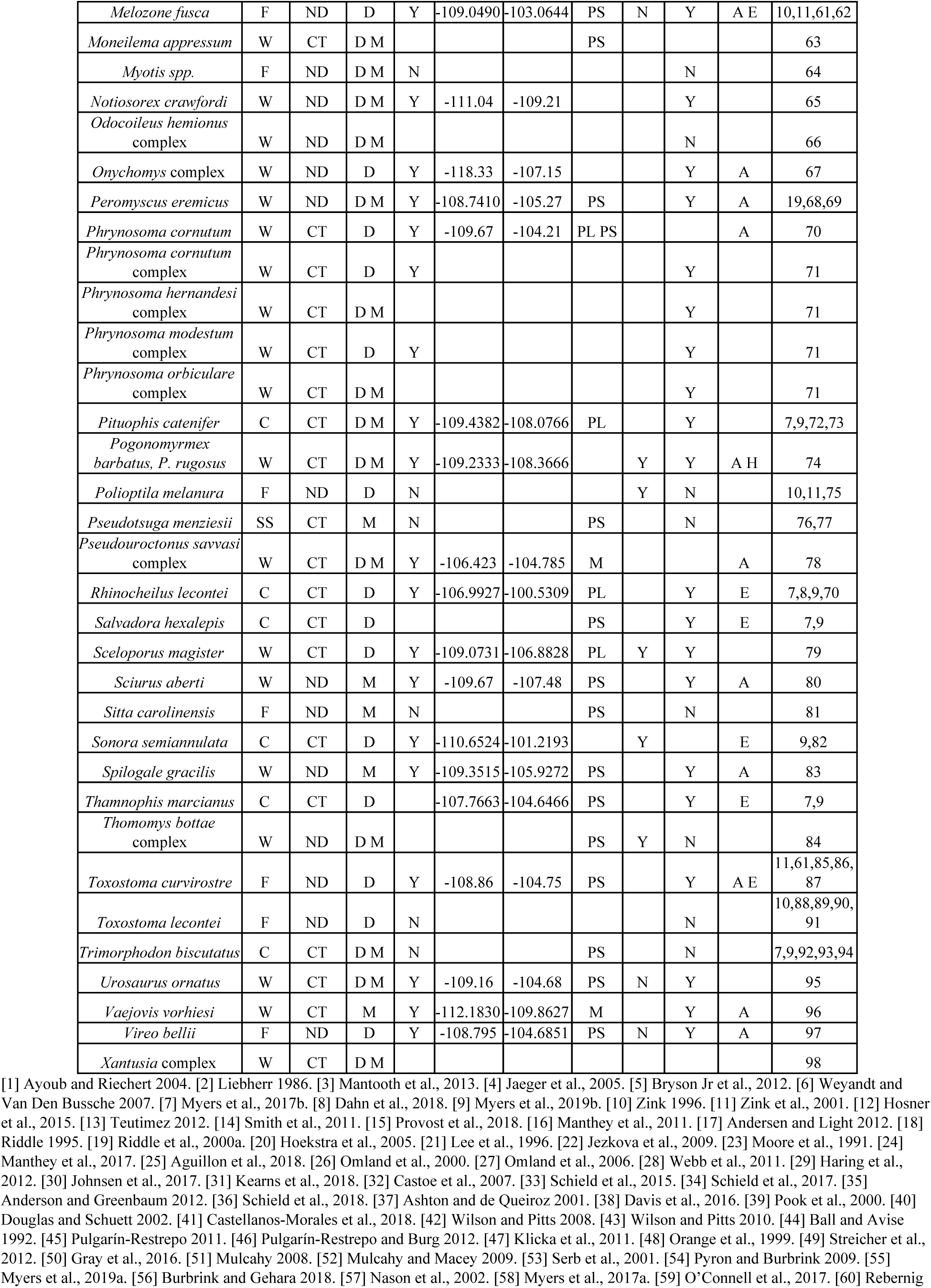
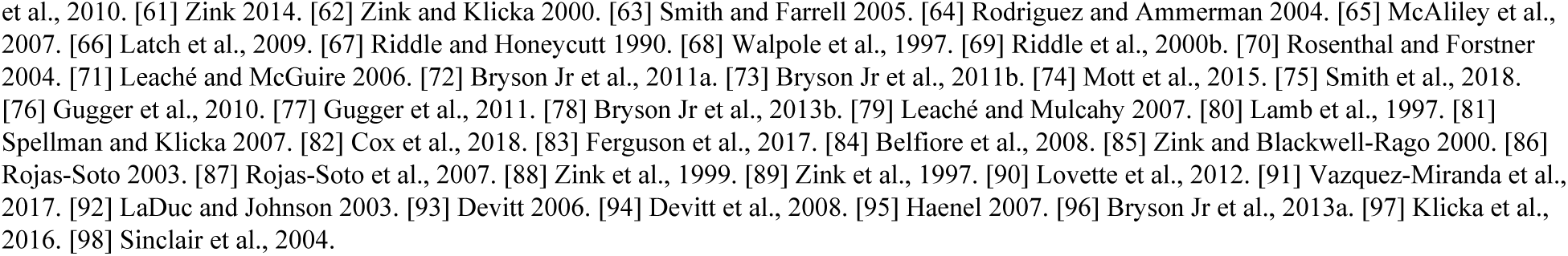
Synthesis of previously published phylogeographic studies. Loco = locomotion types (W= walking, J = jumping, F = flying, C = crawling, SM = swimming, SS = sessile). Therm = thermoregulation types (CT = ectotherm, DC = endotherm). Str = structure across CFB (N = no, Y = yes). West and East are most extreme longitudes of contact zones. Time = divergence times (M = Miocene, PL = Pliocene, PS = Pleistocene). GF = gene flow (N = no, Y = yes). Mono = monophyletic or not (N= no, Y = yes). Mode = mode of speciation (A = allopatry, Ecological, S = sexual selection, H = hybridization, P = polyploidy). See footnote for reference numbers.

**Table S1.4:**
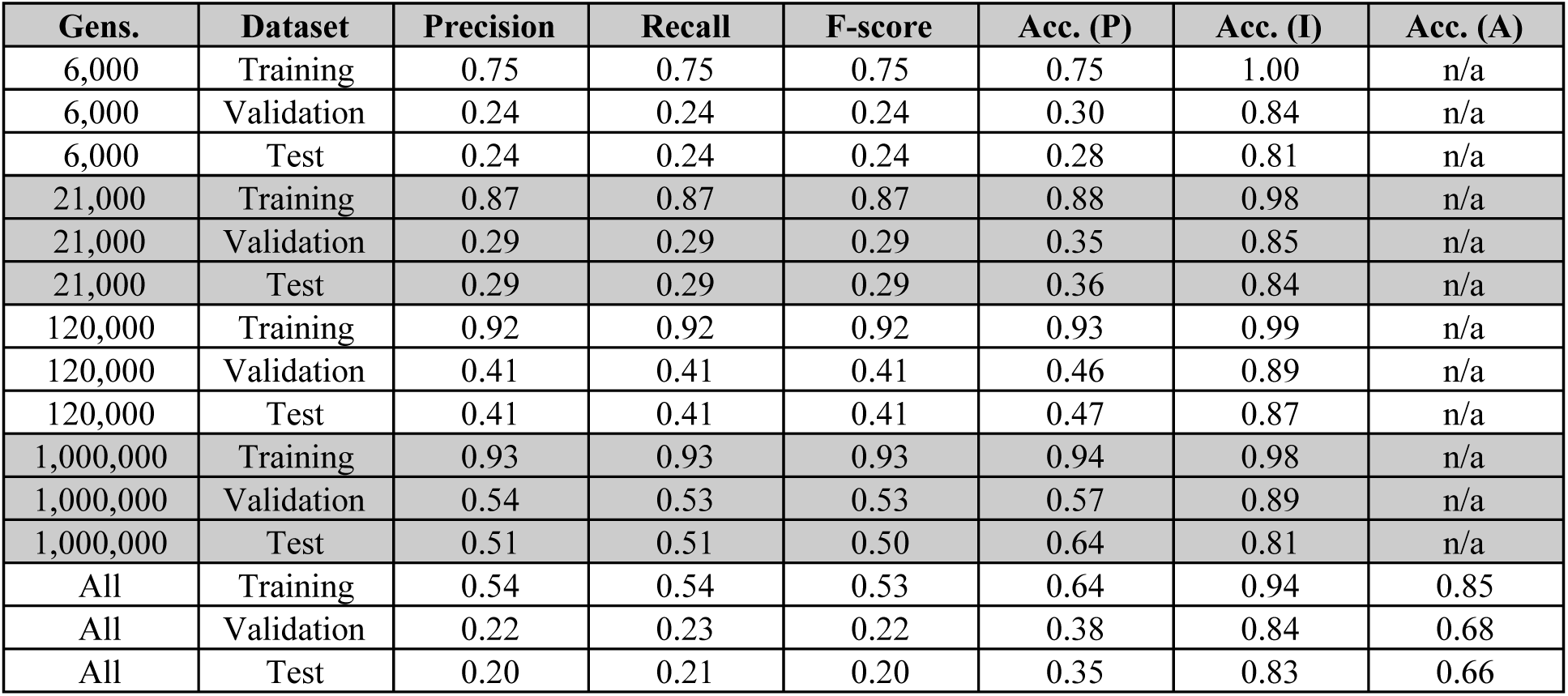
Model results for all 33 statistics. Accuracy scores are calculated for phylogeographic structure alone (P), IBD alone (I), and ages alone (A) when applicable. “Gens.” =generations. “Acc.” =accuracy.

| **Gens.** | **Dataset** | **Precision** | **Recall** | **F-score** | **Acc. (P)** | **Acc. (I)** | **Acc. (A)** |
| --- | --- | --- | --- | --- | --- | --- | --- |
| 6,000 | Training | 0.75 | 0.75 | 0.75 | 0.75 | 1.00 | n/a |
| 6,000 | Validation | 0.24 | 0.24 | 0.24 | 0.30 | 0.84 | n/a |
| 6,000 | Test | 0.24 | 0.24 | 0.24 | 0.28 | 0.81 | n/a |
| 21,000 | Training | 0.87 | 0.87 | 0.87 | 0.88 | 0.98 | n/a |
| 21,000 | Validation | 0.29 | 0.29 | 0.29 | 0.35 | 0.85 | n/a |
| 21,000 | Test | 0.29 | 0.29 | 0.29 | 0.36 | 0.84 | n/a |
| 120,000 | Training | 0.92 | 0.92 | 0.92 | 0.93 | 0.99 | n/a |
| 120,000 | Validation | 0.41 | 0.41 | 0.41 | 0.46 | 0.89 | n/a |
| 120,000 | Test | 0.41 | 0.41 | 0.41 | 0.47 | 0.87 | n/a |
| 1,000,000 | Training | 0.93 | 0.93 | 0.93 | 0.94 | 0.98 | n/a |
| 1,000,000 | Validation | 0.54 | 0.53 | 0.53 | 0.57 | 0.89 | n/a |
| 1,000,000 | Test | 0.51 | 0.51 | 0.50 | 0.64 | 0.81 | n/a |
| All | Training | 0.54 | 0.54 | 0.53 | 0.64 | 0.94 | 0.85 |
| All | Validation | 0.22 | 0.23 | 0.22 | 0.38 | 0.84 | 0.68 |
| All | Test | 0.20 | 0.21 | 0.20 | 0.35 | 0.83 | 0.66 |

**Figure S1.1:**
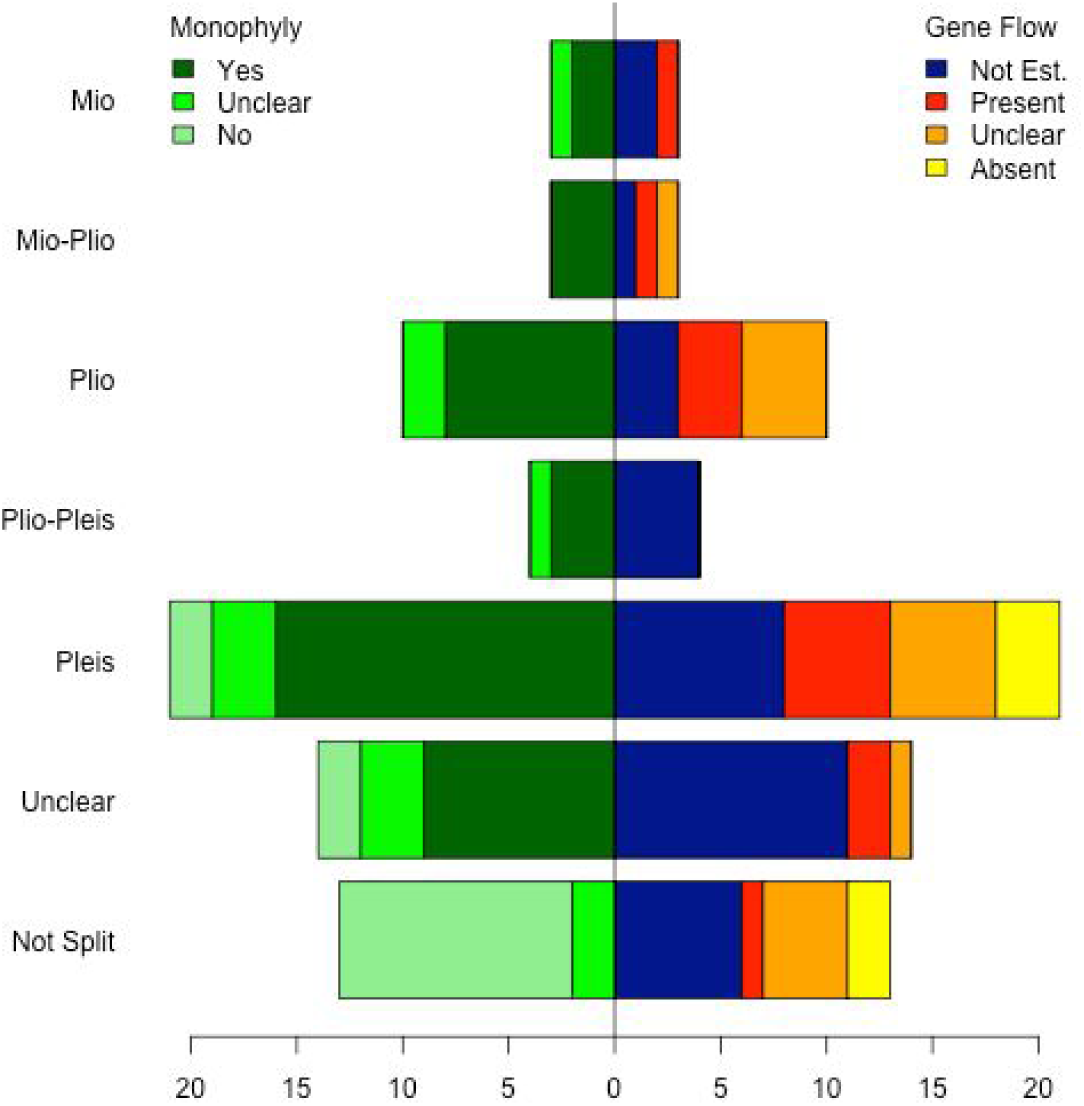
Most of the studied taxa across the Cochise Filter Barrier split during the Pleistocene and did not have gene flow assessed. Monophyly and gene flow assessment varies across taxa and across time. Y-axis shows the timing of the split across the Cochise Filter Barrier (if present). X-axis shows the number of taxa that fall into each category. Left side of the figure shows whether monophyly across the CFB in the taxa is present (dark green), unclear (bright green), or absent (pale green). Right side of the figure shows gene flow. Colors indicate if gene flow is unestimated (dark blue), present (red), unclear (orange), or absent (yellow).

**Figure S1.2:**
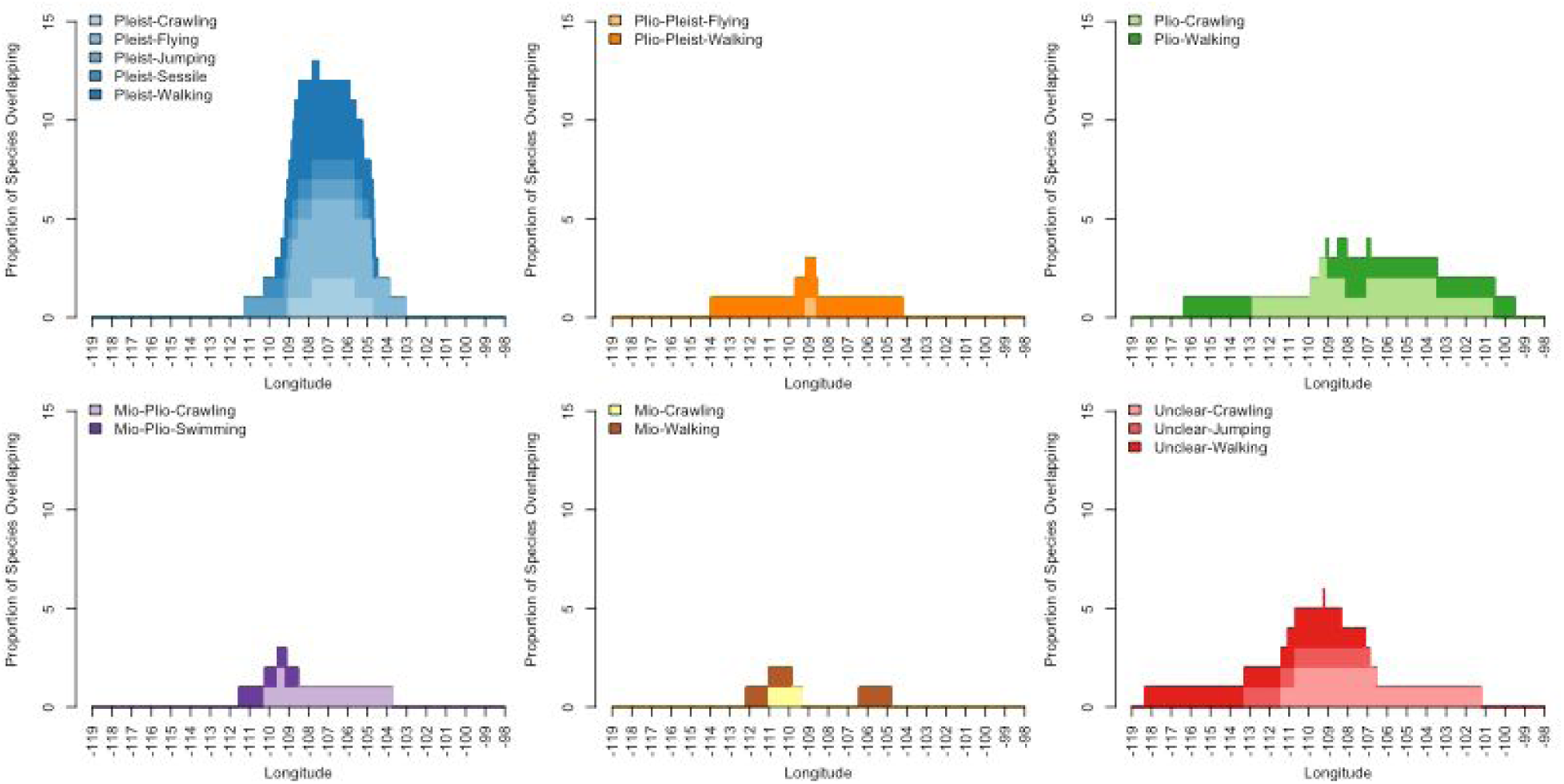
Taxa that diverge in the Pliocene show a wider CFB than taxa that diverge in other time frames. Colors, hues, and shades as in Figure 3. Top panels from left to right: Pleistocene, Pliocene, and Plio-Pleistocene. Bottom panels from left to right: Mio-Pliocene, Unknown, and Miocene.

**Figure S1.3:**
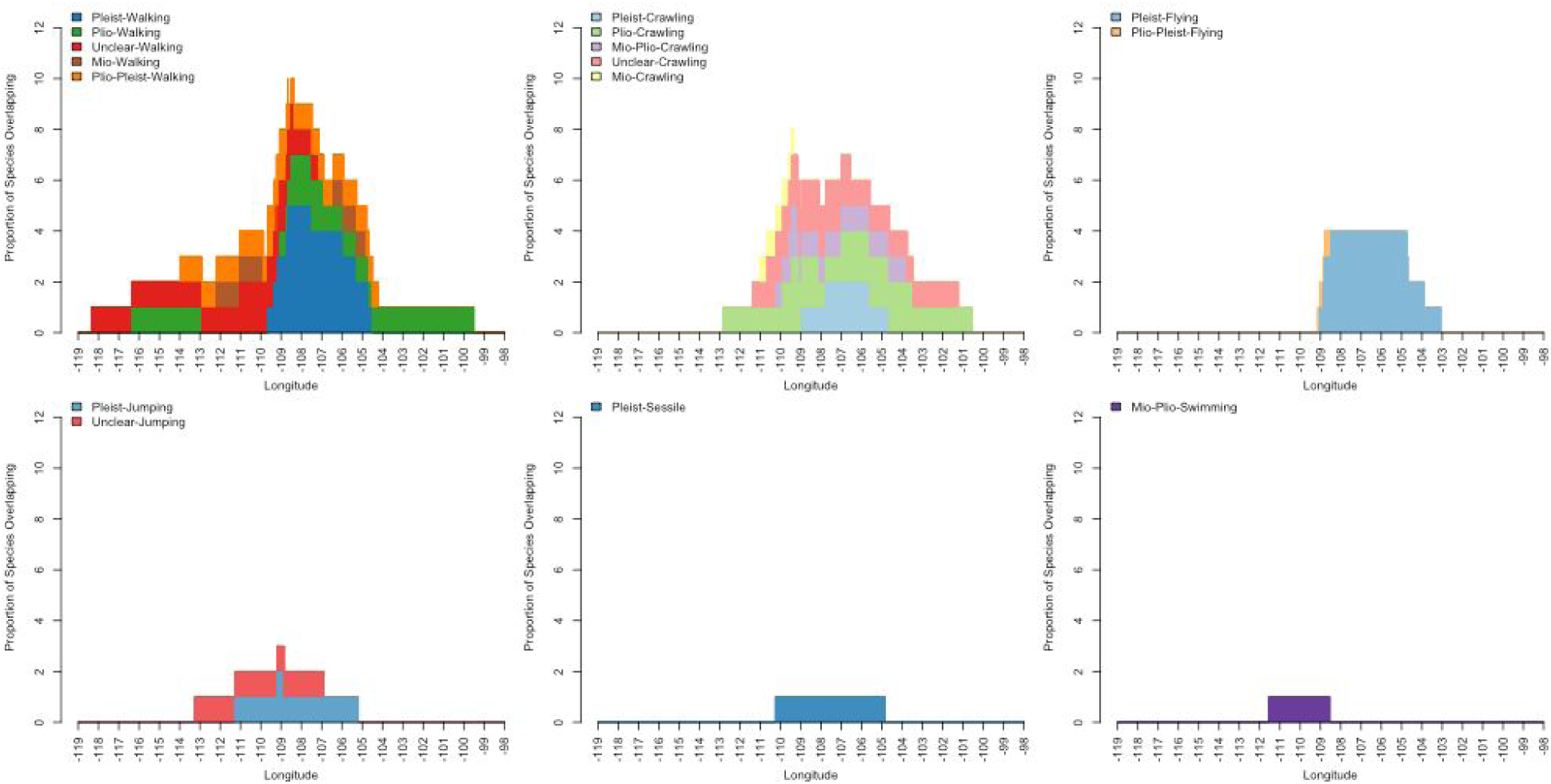
Taxa that walk show a wider CFB than taxa that crawl or fly. Colors, hues, and shades as in Figure 3. Top panels from left to right: crawling, flying, and jumping. Bottom panels from left to right: sessile, walking, and swimming.

**Figure S1.4.**
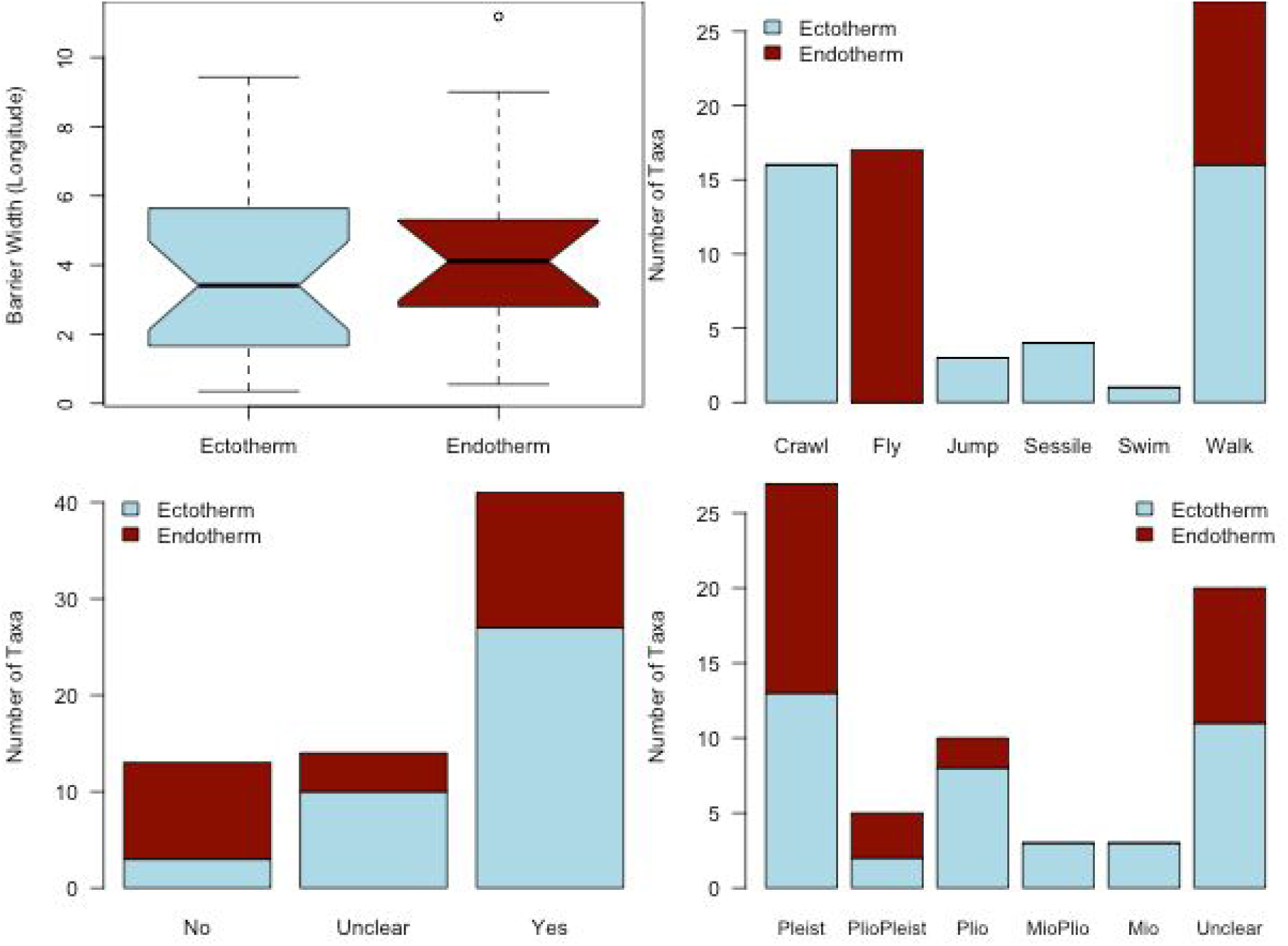
Endothermic species (blue-grey) show more consistent contact zone widths, proportionately less phylogeographic structure, and younger divergence times than ectothermic species (dark red). Top left: boxplot showing the width of the barrier’s contact zone (longitude) between ecto- and endo-thermic species. Top right: number of species with different locomotion types that are ecto- or endo-thermic. Bottom left: the number of taxa that show presence, absence, or ambiguous phylogeographic structure (see legend colors) for endo- and ecto-thermic species. Bottom right: divergence epochs for taxa with phylogeographic structure colored by endo- and ecto-thermic species.

**Figure S1.5:**
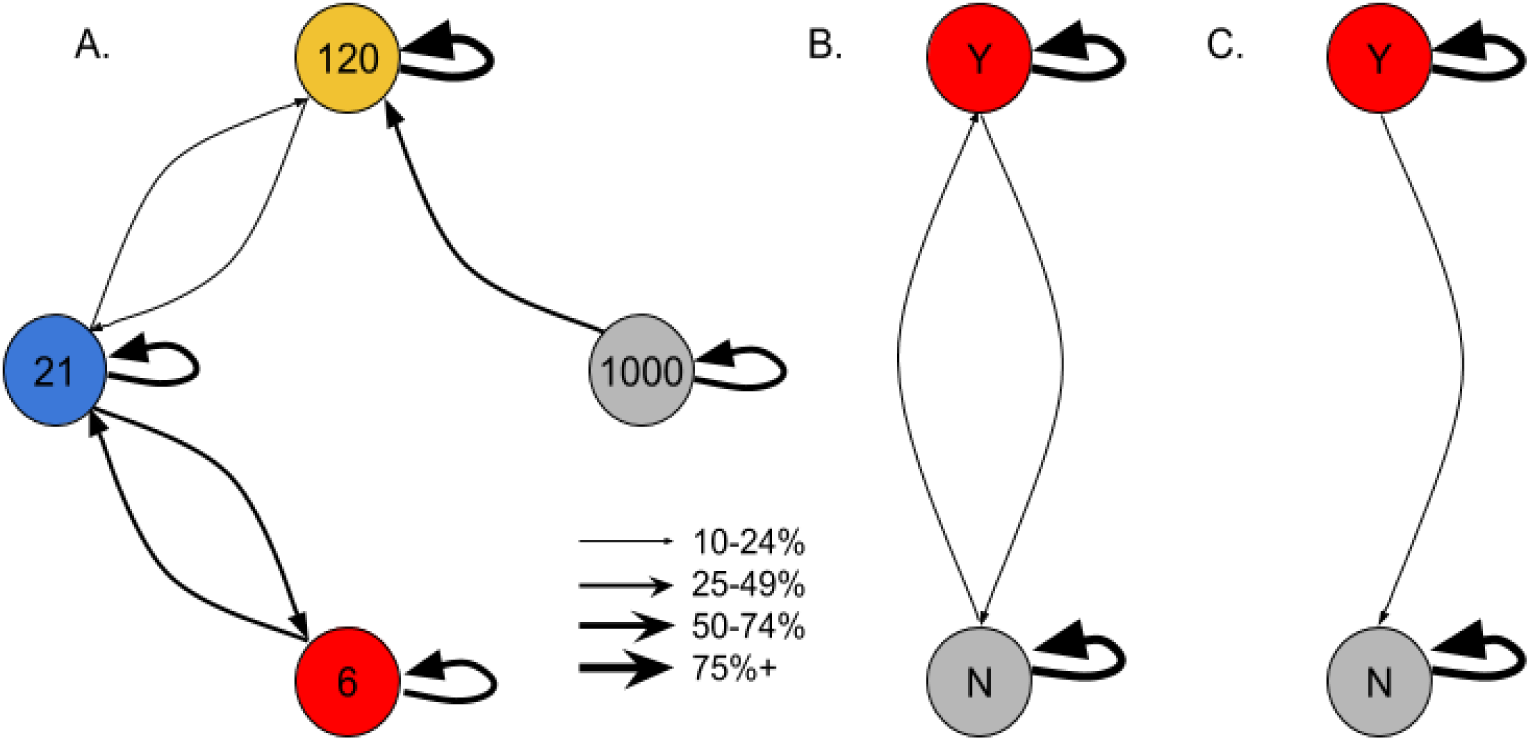
Generations since divergence and presence/absence of IBD are easy to classify. Arrows indicate model assignments. Arrows as in Figure 6. A) Confusion between different ages, 6,000 generations (“6”) through 1,000,000 generations (“1000”). B) Confusion between IBD present (“Y”) and absent (“N”) for 6,000-120,000 generations. C) As in B for 1,000,000 generations.

**Figure S1.6:**
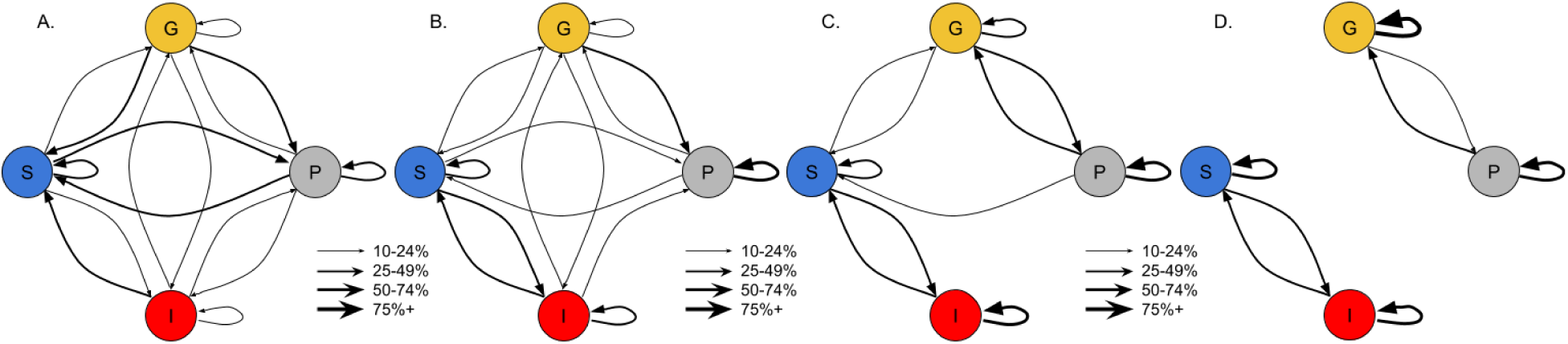
Phylogeographic structure is easy to classify after a long time since divergence, but not after a short time. Arrows and demographic prefixes (“P”, “G”, “I”, and “S”) as in Figure 6. A) 6,000 generations. B) 21,000 generations. C) 120,000 generations. D) 1,000,000 generations.

